# Hybrid Protocells based on Coacervate-Templated Fatty Acid Vesicles combine Improved Membrane Stability with Functional Interior Protocytoplasm

**DOI:** 10.1101/2024.08.06.606659

**Authors:** Jessica Lee, Fatma P. Cakmak, Richard Booth, Christine D. Keating

## Abstract

Prebiotically-plausible compartmentalization mechanisms include membrane vesicles formed by amphiphile self-assembly and coacervate droplets formed by liquid-liquid phase separation. Both types of structures form spontaneously and can be related to cellular compartmentalization motifs in today’s living cells. As prebiotic compartments, they have complementary capabilities, with coacervates offering excellent solute accumulation and membranes providing superior boundaries. Herein, we describe protocell models constructed by spontaneous encapsulation of coacervate droplets by mixed fatty acid/phospholipid and by purely fatty acid membranes. Coacervate-supported membranes formed over a range of coacervate and lipid compositions, with membrane properties impacted by charge-charge interactions between coacervates and membranes. Vesicles formed by coacervate-templated membrane assembly exhibited profoundly different permeability than traditional fatty acid or blended fatty acid/phospholipid membranes without coacervate interiors, particularly in the presence of Mg^2+^ ions. While fatty acid and blended membrane vesicles were disrupted by addition of 25 mM MgCl_2_, the corresponding coacervate-supported membranes remained intact and impermeable to externally-added solutes even in the presence of MgCl_2_. With the more robust membrane, fluorescein diacetate (FDA) hydrolysis, which is commonly used for cell viability assays, could be performed inside the protocell model due to the simple diffusion of FDA and then following with the coacervate-mediated abiotic hydrolysis to fluorescein.

## 1. Introduction

Compartmentalization is crucial to how protocellular systems are thought to have formed on the early earth, providing a means of (co)localizing molecular components, facilitating reactions, and defining individuals upon which natural selection could operate.^1–3^ Potential compartmentalization mechanisms include coacervate droplets formed by liquid-liquid phase separation and membrane vesicles formed by amphiphile self-assembly.^4–8^ Coacervates can accumulate molecular solutes such as nucleic acids^9–15^, accumulating them to high local concentrations in their interior while depleting these solutes from the bulk continuous phase. However, these droplets undergo coalescence upon contact and the absence of a boundary membrane limits resistance to solute entry/egress.^4, 16^ In contrast, membrane vesicles formed from single-chain amphiphiles such as fatty acids provide such a boundary while lacking a general mechanism for solute accumulation from dilute solution.

Permselective molecular transport is a defining characteristic of lipid membranes and an important benefit of their use as protocellular compartments. For example, lipid vesicles can compartmentalize RNA, and host RNA catalysis and self-replication.^17–19^ However, the addition of single-chain fatty acids, which are considered more prebiotically-plausible,^20, 21^ to phospholipid membranes increases membrane permeability to solutes,^22–25^ with greater permeability for higher amounts of fatty acid in the membranes. Higher membrane permeability could be beneficial at early stages of prebiotic compartmentalization when membrane transporters such as transmembrane proteins would not be available. However, fatty acid or fatty acid/phospholipid vesicles are susceptible to disruption by Mg^2+^,^18, 19^ which interacts with their headgroups. This is a disadvantage for functional RNA compartmentalization and the “RNA World” hypothesis since Mg^2+^ commonly participates in RNA folding and catalysis.^26, 27^ Additionally, passive loading of solutes such as RNAs into lipid vesicles by encapsulation during formation provides a relatively low molecule encapsulation efficiency, with internal solute concentrations approximating external concentrations and hence usually only a tiny fraction of the total solutes located inside.^28–32^ Therefore, hybrid structures comprising lipid membrane-bounded coacervate droplets are a promising new approach to combine the benefits and address the drawbacks of these compartment types.

Today’s cells are themselves compartments and are further organized into organelles based on lipid membranes and liquid-liquid phase separation.^33,34^ Increasingly, membranes and liquid phases are found to interact as part of biological functions that include transmembrane signaling, immune cell activation, and actin assembly.^35–37^ Membrane-condensate interactions drive liquid-liquid phase separation, membrane organization, and condensate size.^35, 37–41^ Yet, it remains unclear what factors are most critical in droplet–membrane interactions. Simple model systems have been designed to provide insight into how these two organelle-forming assemblies interact with each other.^5, 42^ Coacervates or biocondensates have been introduced to giant unilamellar lipid vesicles as models of organelles^43–45^; recently, the self-assembly of lipid membranes at the surface of coacervate droplets via several methods has been used to construct protocell and synthetic cell models that take advantage of combining a membrane boundary and macromolecularly crowded interior “cytoplasm”.^20, 21, 46–50^ The permeability properties of these coacervate-supported membranes are only just beginning to be studied, but so far all reported permeability tests have shown reduced barrier function for coacervate-supported membranes as compared with traditional vesicle membranes.^46, 47^ This can be beneficial, for example when membranes selectively pass small molecules that can serve as reaction substrates while retaining larger molecules such as enzymes in the interior lumen,^47^ or when populations of coacervate core vesicles with heterogeneous membrane permeability offer the possibility of selective pressure.^48^ At the same time, enhanced permeability suggests disruption of the expected molecular organization within the lipid bilayers. Is such disruption an unavoidable consequence of the lipid-coacervate interactions that enable formation of coacervate-supported membranes? What impact does contact with the coacervate interior have on membrane stability to challenges such as externally added Mg^2+^? Much remains to be learned about the mechanisms of membrane formation, the structures and properties of the resulting membranes, and how to predict and design coacervate-supported membranes.

Here, we examined the formation and properties of coacervate-supported membranes based on single-chain fatty acids and phospholipid/fatty acid mixtures, with an emphasis on membrane permeability and robustness to externally-added Mg^2+^. Coacervate droplets prepared from a structurally simple synthetic polyamine (polyallylamine hydrochloride, PAH) and adenosine diphosphate (ADP) were used as templates for lipid self-assembly via a gentle hydration approach. Membranes were assembled from oleic acid alone or mixtures of oleic acid with 1-palmitoyl-2-oleoyl-glycero-3-phosphocholine (POPC). The importance of the charge-charge interaction was evaluated by independently varying the membrane and coacervate surface charges. Membrane permeability to several fluorescently-labeled solutes was compared for mixed and pure fatty acid membranes with and without coacervate cores. We also examined the impact of 25 and 50 mM added Mg^2+^ on stability and permeability of these coacervate-supported membranes. We found that the coacervate-supported membranes exhibit profoundly different permeability than traditional, unsupported fatty acid or blended fatty acid/phospholipid membranes, particularly in the presence of Mg^2+^. The coacervate-supported membranes were able to block solute entry to the coacervate interiors, even in the presence of Mg^2+^. The unexpectedly low solute permeability and better divalent metal stability of these fatty acid membranes suggest that combining prebiotically-plausible single-chain amphiphile assembly with coacervate proto-cytoplasms improves the barrier function and robustness of fatty acid membranes. Finally, we exposed the membrane-coated coacervate hybrid protocell models to fluorescein diacetate (FDA). This neutral small molecule was able to pass the coacervate-supported membranes, whereupon the amine-rich coacervate interior catalyzed ester hydrolysis to produce a green fluorescent product, fluorescein. The negative charge on fluorescein greatly slows its exit from the protocell interior, causing it to accumulate. This very simple abiotic reaction echoes the way living cells process this common cell viability test reagent, illustrating how combined function from both the reactive coacervate interior and the semipermeable membrane can begin to suggest steps towards “metabolism”-like reactions in the hybrid protocells.

## 2. Results and discussion

**Figure 1.**
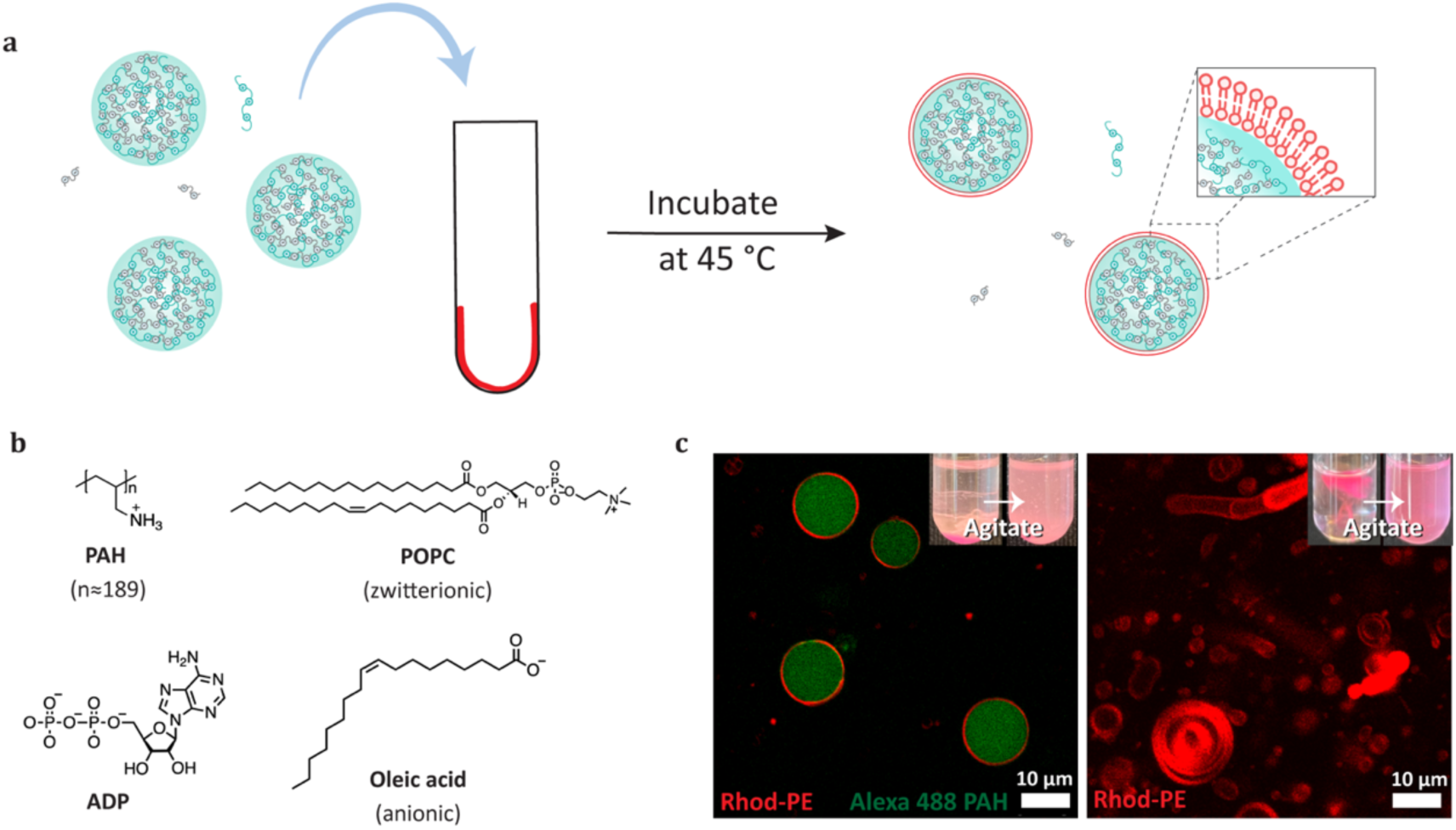
Formation of coacervate-supported membranes. (A) Scheme of the formation of coacervate-supported membrane via gentle hydration method using coacervate solution. (B) Molecular structures of polyelectrolytes, poly(allylamine hydrochloride) (PAH) and adenosine-5’-diphosphate (ADP), used for making coacervates and lipids, POPC and oleic acid, used for lipid assembly. (C) Confocal fluorescence and microscopy images of coacervate-supported membranes and vesicles made with POPC 49.95 mol%, oleic acid 50 mol%, and Rhodamine labeled PE 0.05% in the presence and absence of PAH:ADP coacervates (20 mM charge concentration each) with 250 mM Tris at pH 8.7. Fluorescence images have been contrast-adjusted and false-colored for ease of visualization: green and red indicate Alexa 488 labeled PAH and Rhod-PE respectively.

Coacervate-supported membranes (CSM) were formed by a modified gentle hydration approach ^48^ (Figure 1A). We hydrated phospholipid/fatty acid blended lipid films by adding a coacervate suspension composed of PAH and ADP (Figure 1B), where PAH is a chemically simple model polycation and ADP was chosen as an oppositely-charged small molecule that is prebiotically plausible and important for extant biology. We observed that the coacervate droplet interface served as a template for lipid self-assembly (Figure 1C). Many of the coacervate-templated membrane structures observed by confocal fluorescence and fluorescence polarization appeared to be thin and relatively uniform (Supplementary Figure 1A), suggesting oligolamellar or bilayer membranes as opposed to the thicker multilayers observed by Tang et al^46^ for OA assembled from solution. These CSM appear more similar to the bilayer membranes found previously for diacyl phospholipid assembly formed by hydrating dried lipid films in coacervate-containing media.^48^ Polarization of emission from Lissamine rhodamine b labeled at the DOPE lipid in the membrane indicates molecular orientation consistent with lipid ordering within the membrane.^51, 52^ We observed higher polarization from POPC+OA CSM than OA-CSM (Supplementary Figure 1B and C), which is consistent with the expected higher mobility of lipids in single chain fatty acid membranes. In contrast to the relatively thin, similarly-sized, spherical CSMs, lipid assemblies from traditional gentle hydration of these lipid mixtures were relatively heterogeneous in sizes, shapes, and lamellarity, as expected for this giant vesicle preparation method.^53^ We noticed that the yield of OA-CSMs was relatively lower than POPC+OA-CSMs (Supplementary Figure 2). This could be due to the higher solubility of the fatty acid as compared to phospholipid.

**Figure 2.**
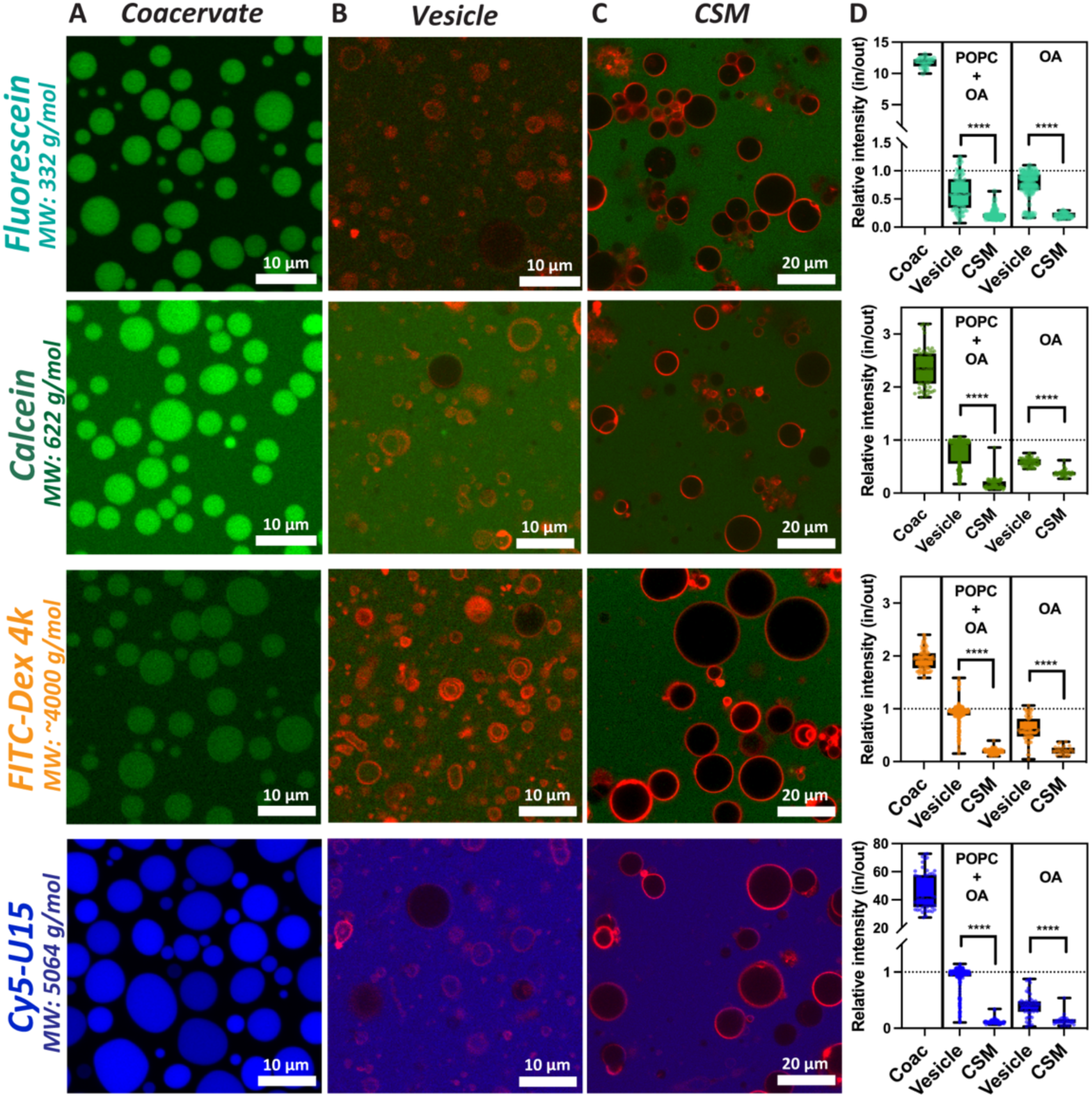
Coacervate-supported phospholipid/fatty acid membranes are impermeable to fluorescein, calcein, dextran 4kDa, and oligoRNA. Permeability test of POPC+OA (each 50 mol%)-coacervate supported membrane (CSM) using various solutes. Addition of solutes including anionic molecules (Fluorescein and Calcein, green in the first and second column, correspondingly), labeled neutral molecule (FITC-dextran 4k, green in the second column), and labeled RNA (Cy5-U15, blue) to (A) PAH:ADP coacervates, (B) POPC+OA-vesicles without coacervates, and (C) POPC+OA-CSM. Red fluorescence indicates Rhod-PE. All images were taken 20 mins after the addition of solutes, except for Cy5-U15 added to coacervates was taken 1 day after addition to allow for slower equilibration due to its stronger partitioning. (D) Relative intensity (in/out) of three solutes in coacervate, vesicle, and CSM after 20 mins. All vesicle and CSM data for three solutes and for two kinds of membrane are statistically different (p-values are < 0.0001) except fluorescein added to OA-CSM. Fluorescence images have been contrast-adjusted and false-colored for ease of visualization Intensity ratios were calculated from raw data after subtraction of dye-free background signal from *I_inside_* and *I_outside_*.

### 2.1. Coacervate-supported fatty acid-phospholipid blended and fatty acid membranes resist solute entry

One advantage of having a membrane is to provide a physical barrier in order to control access of molecules to/from the interior volume. However, oleic acid and oleic acid/phospholipid blended membranes are known to exhibit solute leakage in GUVs^22–25^. To compare the POPC+OA and OA lipid membranes prepared in the presence/absence of coacervates, we tested the permeability using several different solutes, including two anionic small dyes (Fluorescein and Calcein), a neutral small dye (Nile Red), a cationic small dye (Propidium iodide, PI), a labeled neutral molecule (FITC-Dextran 4k), and a labeled biomolecule (Cy5-U15, RNA). Since coacervates themselves can sequester or exclude different solutes, we made sure that all solutes tested here were able to accumulate inside coacervates, with relative fluorescence intensities (in/out) >1. Each of the six solutes met this criterion. Each of the remaining fluorescent solutes were then added to systems of coacervates only, vesicles only (without coacervates), and coacervate-supported membranes. Nile Red and PI both accumulated in or at the membrane, making assessment of permeability difficult, therefore these dyes were excluded from further study (see Supplementary Figure 3 and Supporting Discussion).

For the remaining four solutes (fluorescein, calcein, FITC-dextran 4k, and Cy5-U15 RNA), images were collected for at least 35 structures for statistical comparison, and fluorescence intensity ratios determined as the ratio of fluorescence intensity inside to outside 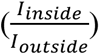. Intensity ratios were calculated from raw data after subtraction of background signal, which was obtained by the sample with no fluorescent solute added. For coacervate samples, these ratios represent apparent partitioning coefficients. We note that fluorescence intensities may be impacted by their local environment, particularly inside coacervates, making confident quantification difficult particularly as medium-appropriate calibration data could not be acquired for the dense coacervate phase. Although fluorescence intensity is an imperfect measure of solute concentration, the appearance of a fluorescent solute in the interior of a vesicle or CSM clearly indicates its ability to permeate the boundary membrane. Fluorescence intensity values further provide insight into the variability across structures within a population of vesicles, coacervates, or CSM.

Figure 2 summarizes the results. Coacervates alone accumulated all solutes to concentrations higher than in the surrounding solution, with average fluorescence intensity ratios of ∼2x higher inside for calcein and FITC-dextran 4k, ∼12x higher inside for fluorescein, and ∼46x higher inside for Cy5-U15 RNA (Figure 2A, D). The much higher uptake of labeled RNA oligomer is expected based on its multiple negative charges and ability to compete with ADP for binding to the polyamines in the coacervate.^9, 10, 54^ POPC+OA vesicles had, on average, nearly equal fluorescence intensities inside and outside, indicating that the labeled solutes were able to permeate through the POPC+OA membranes and approach equilibrium concentrations (Figure 2B, D). In the absence of a coacervate interior, there is no driving force for solute accumulation in the vesicle interiors, and an intensity ratio in/out of 1.0 indicates complete equilibration of solute across the membrane. Our observation of intensity ratios ∼ 1 for POPC+OA vesicles are generally consistent with literature reports of solute leakage from vesicles made from fatty acid-containing membranes.^24^ In contrast, the coacervate-supported POPC+OA membranes effectively excluded all three solutes, despite the much greater driving force for their accumulation within the coacervate interior as compared to the vesicle lumens (Figure 2C). Average intensity ratios for solutes in the POPC+OA-CSM samples were ∼0.2 for calcein and FITC-Dex 4kDa, and ∼0.1 for Cy5-U15 (Figure 2D). The lower value for Cy5-U15 despite its higher partitioning preference for the coacervates is consistent with its greater negative charge and larger MW reducing membrane permeability as compared to the other solutes.

We considered the possibility that our relatively high pH of 8.7 could be playing some role in our results. This pH is near the pKa of OA in lipid membranes^55–57^ and was chosen to favor OA membrane assembly by enabling hydrogen bonding between protonated and deprotonated headgroups^57^. Previous literature has shown that the introduction of increasing amounts of oleic acid to POPC membranes caused leakage at pH 8.^24^ Therefore, we also performed the permeability test using calcein at pH 8.0 and 7.4 (physiological pH). We observed similarly strong barrier function for the coacervate-supported fatty acid/phospholipid membranes under these conditions (Supplementary Figure 4).

We also tested OA-only membrane vesicles and OA-only CSM, and found similar trends (Figure 3D). OA-only CSM had statistically significant lower average intensity ratios than vesicles (Supplementary Table 1), similar to that of the POPC+OA membranes, and were more effective in preventing accumulation of calcein, FITC-Dex 4k, or Cy5-U15 RNA in their coacervate cores. We noticed that OA-only vesicles were themselves better at resisting solute entry than the POPC-OA blended vesicles in this experiment, specifically for calcein and Cy5-U15 RNA. This was unexpected since OA vesicles are known to be “leaky”^24^, and could be the result of electrostatic repulsion between these negatively charged solutes and the negatively charged lipid membrane. However, when incubated for longer times than the 20 minutes used for the experiments of Figure 3, pure OA vesicles were unable to prevent entry of the negatively-charged dye, calcein, to the vesicle lumen: by 24 hrs it was nearly equilibrated, in contrast to the larger FITC-Dex 4k, which remained excluded (Supplementary Figure 5). Therefore, our fatty acid permeability data appears generally consistent with literature^22, 24, 46^.

Some heterogeneity in solute entry was observed between structures within a population (note error bars in Figure 2D). Variability was greatest for the vesicle samples, with some vesicles effectively excluding solutes while others within the same population equilibrated fully with the exterior solute concentration. Differences in lamellarity of the vesicle membranes and CSMs within a population are suggested based on their differences in membrane fluorescence intensity.^58^ We were unable to directly correlate these differences with solute entry in our samples due to additional forms of heterogeneity of the populations (varying diameters meant that some membranes were imaged above or below their equators, some structures are touching or attached to lipid aggregates, etc.). However, overlaid confocal fluorescence images such as those shown in Figure 2 do not suggest a strong correlation between membrane thickness (as intensity in the red channel) and solute entry (as intensity in the green or blue channels), and dark interiors are seen for all of the CSMs (Figure 2C, D).

Histograms comparing intensity ratios (for solute in/out) for populations of coacervates, POPC+OA-vesicles, and POPC+OA-coacervate-supported membranes are shown in Supplementary Figure 6. The percent of vesicles showing solute penetration through the membrane (defined here as relative fluorescence intensities in/out > 0.5) for fluorescein, calcein, FITC-Dex 4k, and Cy5-U15 for vesicles are 63.1% (out of 130), 73.9% (out of 352 vesicles), 84.8% (out of 374 vesicles), and 83.4% (out of 308 vesicles) correspondingly. On the other hand, close to or fewer than 1% of CSM showed solute penetration out of more than 135 coacervate-supported vesicles (Supplementary Table 2). For POPC+OA-vesicles (without coacervate interior), the relative fluorescence intensities for labeled solutes inside vs outside are widely distributed from << 1 to ∼ 1, indicating variability in membrane structure across the population. In contrast, the POPC+OA-CSM in general had smaller internal fluorescence intensity for all three solutes, indicating that contact with the coacervate interiors alters the lipid membrane’s barrier property and prevents solute entry. Superior barrier function for the coacervate-supported blended fatty acid/phospholipid membranes as compared to vesicles of the same lipid composition was notable, particularly since previous reports from our lab^48^ and others^46, 47^ showed the opposite behavior, finding increased solute permeability for coacervate-supported membranes based on other coacervate/lipid combinations. In order to gain a more systematic understanding of coacervate-membrane interactions, we next focused on varying the charge of the coacervate and membrane components while maintaining the same molecular identities.

**Figure 3.**
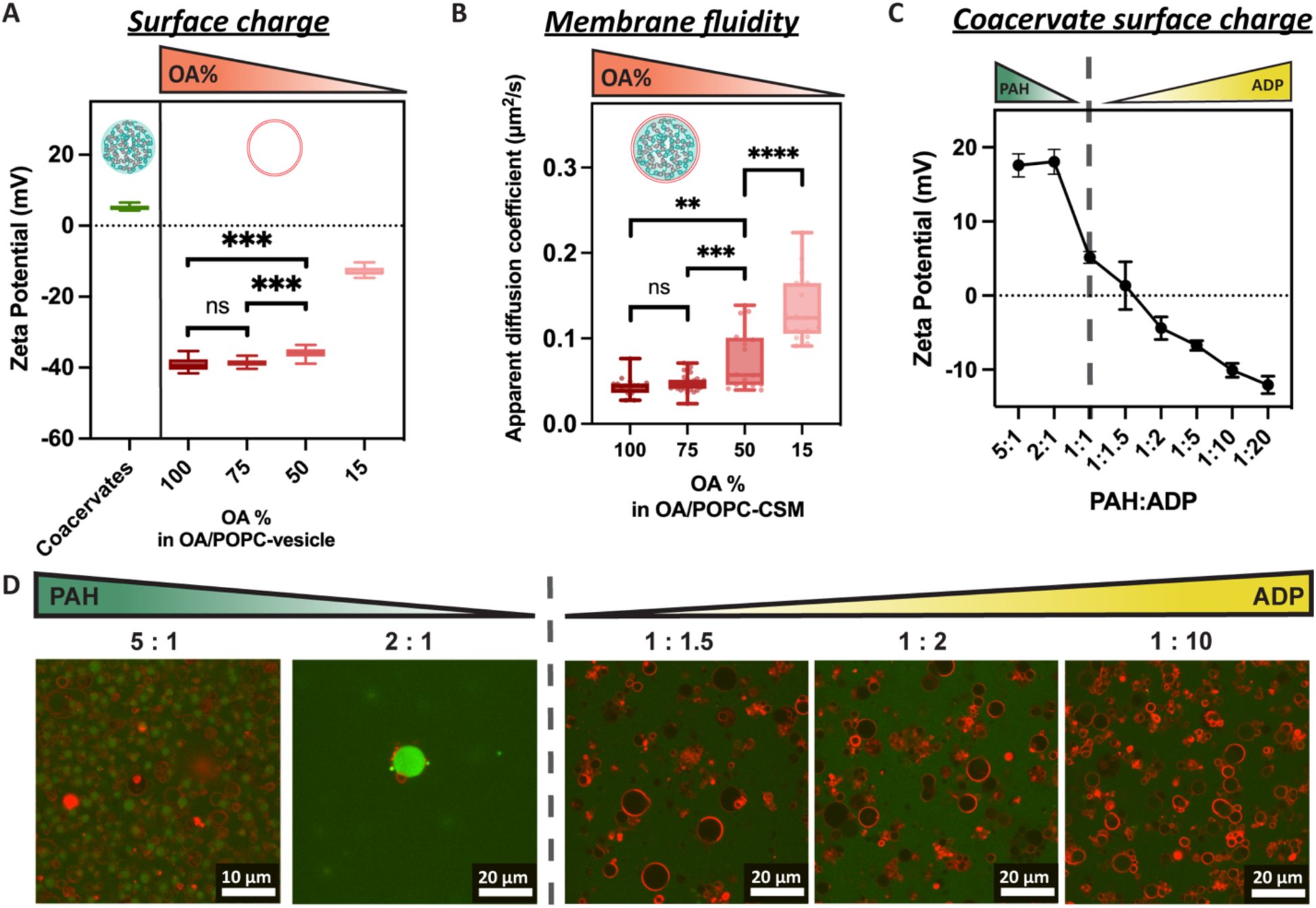
Charge-charge interactions between lipid membrane and coacervate droplet surfaces influence assembly of coacervate-supported membranes. (A) Zeta potential measurements of 1:1 (PAH:ADP) coacervates and of lipid vesicles as a function of the amount of negatively charged lipid (oleic acid) in the membrane, the balance of which is composed of zwitterionic POPC. (B) Apparent diffusion coefficients of labeled lipid molecules within CSM formed from lipid mixtures having varying ratios of OA:POPC. (C) Zeta potential measurements of coacervates with various charge ratio. (D) Images showing lipid assemblies and calcein distribution for coacervate-supported membranes prepared using coacervates with unequal charge ratios; calcein was added externally as a test of membrane permeability. Fluorescence images have been contrast-adjusted and false-colored for ease of visualization (red and green channel indicates Rh-PE, and calcein fluorescence, respectively).

### 2.2. Membrane-coacervate interactions influence formation and properties of coacervate-supported membranes

We hypothesized that electrostatic interactions between the coacervate surface and lipids were playing a pivotal role in membrane assembly. To test this, we varied the compositions of the lipid membrane and coacervates separately. First, we varied the relative amounts of negatively-charged OA and zwitterionic POPC used to form membranes (Figure 3A). The 1:1 PAH:ADP coacervate system we used has a slightly positive zeta potential (ca. + 5 mV), consistent with an expectation that the longer polyamine molecules are more effectively compartmentalized as compared to the shorter ADP molecules.^9, 10^ All of the vesicles tested here had a negative zeta potential, as expected for mixtures of negatively-charged (OA) and zwitterionic (PC) headgroups. The zeta potential became less negative for membranes having decreased amounts of oleic acid (from around -40 for pure OA to -15 mV for 15% OA).

To understand the interactions between coacervate and lipid membrane, we prepared CSM with each of the OA/POPC compositions from Figure 3A (images can be found in Supplementary Figure 7). For pure POPC composition, we observed lipid aggregates at the coacervate droplets instead of complete lipid membrane. We conducted fluorescence recovery after photobleaching (FRAP) to compare lateral diffusion for a labeled lipid within the membrane and observed slower fluorescence recovery for membranes formed from lipid mixtures with more OA, consistent with stronger interactions between the positively-charged coacervate droplets and negatively-charged lipids in the membrane (Figure 3B). The calculated apparent diffusion coefficients ranged from as low as ∼0.04 μm^2^/s at 100% OA to as high as ∼0.1 μm^2^/s for 15% OA. These values for CSM are all substantially lower than for vesicles lacking the coacervate core (Figure 3B and Supplementary Figure 8), and lowest values were observed for the pure OA membranes. This could be the result of the attractive interactions between coacervate amines and OA’s anionic headgroups slowing lipid diffusion. POPC+OA (50% OA)-vesicles prepared in buffer had apparent diffusion coefficient ∼0.8 μm^2^/s, 10-fold slower than for POPC+OA (50% OA)-CSM (∼0.08 μm^2^/s), which shows the interactions between membrane and coacervate interior as compared to the buffer. Although the PAH and ADP molecules are concentrated inside the coacervates, some concentration exists in the surrounding dilute phase. Therefore, in order to test the effect of PAH and/or ADP molecules from solution interacting with the lipid headgroups to alter membrane properties, we also performed FRAP experiments on vesicles prepared in the dilute supernatant phase of the coacervate system (Supplementary Figure 8). These vesicles prepared in dilute phase had apparent diffusion coefficient of ∼0.5 μm^2^/s, somewhat reduced from vesicles prepared in buffer but still much higher than CSM, further supporting the importance of lipid-coacervate interactions in the properties of CSM samples (Figure 3B and Supplementary Figure 8).

We next examined the impact of changing the relative amounts of PAH and ADP used in forming the coacervates. Zeta potential measurements were used to identify PAH:ADP charge ratios leading to coacervates with more positive, nearly neutral, and negative coacervate droplets (Figure 3C). We then attempted to form POPC+OA (50% OA)-membranes on coacervates with a range of surface charges. The most positive coacervate (5:1 PAH:ADP) did not support CSM formation; these samples instead contained separate populations of coacervate droplets and lipid vesicles (Supplementary Figure 9, leftmost panel). All of the other tested compositions (2:1, 1:1.5, 1:2 and 1:20 PAH:ADP) formed lipid membranes at the coacervate droplet interface (Supplementary Figure 9). However, the CSM formed on 2:1 PAH:ADP coacervates were permeable to calcein, with substantial calcein accumulation in the droplet interiors within 20 minutes (Figure 3D) Although the higher surface charge of these 5:1 and 2:1 PAH:ADP coacervates could be expected to aid assembly of the negatively-charged membranes, poor barrier function of the resulting CSMs suggests that other factors were more important here. We hypothesize that PAH from the dilute phase may be preventing proper membrane assembly by interacting with the lipid headgroups, rendering the polyamine-coated membranes positively-charged, particularly for the 5:1 PAH:ADP case, where substantial excess PAH is present in the dilute phase (Supplementary Figure 10). At 2:1 PAH:ADP, membranes did form at the interface but were unable to prevent calcein from entering, presumably due to membrane defects caused by excess PAH interacting with the lipid headgroups (Supplementary Figure 9). On the other hand, negatively charged coacervates prepared with excess ADP for example 1:2 PAH:ADP formed higher-quality CSMs impermeable to calcein. Even at 1:10 PAH:ADP, CSM could be formed although the encapsulated coacervates were more heterogeneous. The fact that CSMs form when coacervates have excess ADP and are negatively-charged can be rationalized as indicating the membrane’s ability to compete effectively with ADP molecules for binding to the amines of PAH at the interface of the coacervate droplets.

**Figure 4.**
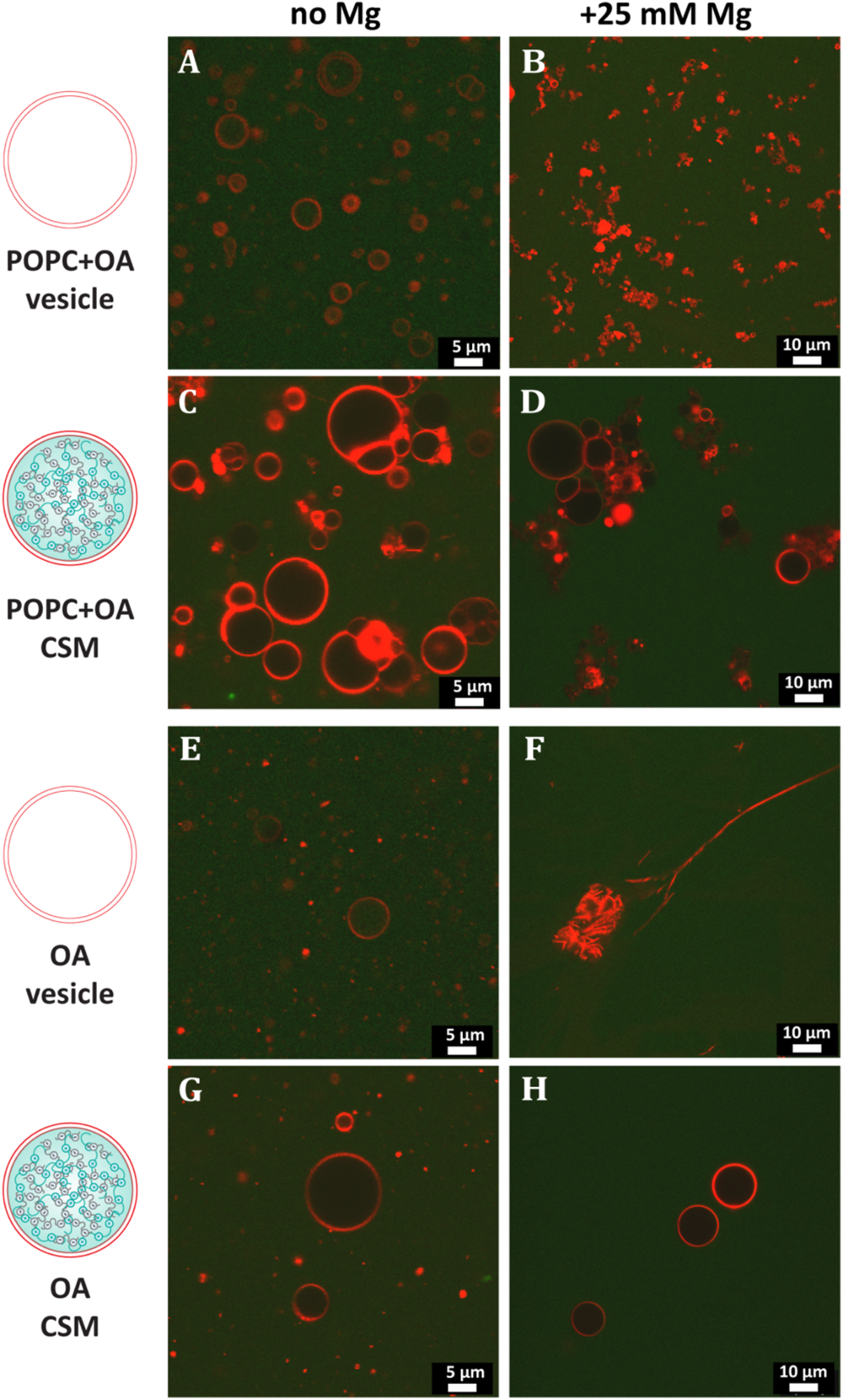
Coacervate-supported membranes retain excellent barrier function after treatment with Mg2+. Confocal fluorescence images for membrane vesicles and coacervate-supported membranes taken 24 hr after addition of FITC-Dex 4k alone (all panels on left) or with 25 mM MgCl_2_ (all panels on right). Blended phospholipid/fatty acid membranes (panels A-D), and fatty acid-only membranes (panels E-H panels), for vesicles (upper panels within each set) and for coacervate supported membranes (lower panels within each set). All coacervates were prepared with 1:1 charge ratio (PAH:ADP). Blended phospholipid/fatty acid membranes had 50 mol % POPC+50 mol % OA, and fatty acid-only membranes had 100% OA. Fluorescence images have been contrast-adjusted and false-colored for ease of visualization (green and red represents FITC-Dex 4k and rhod-PE, respectively).

### 2.3. Coacervate-supported membranes have improved stability to Mg^2+^

The lower permeability of CSM as compared to vesicle membranes without coacervate cores suggested the possibility that CSM might be more stable to unfavorable conditions such as the presence of Mg^2+^. Mg^2+^ disrupts fatty acid membranes by binding to their carboxylate moieties.^24^ This is of particular interest for prebiotic compartmentalization, since proper RNA folding, which is necessary for ribozyme activity, often requires divalent metal cations, most commonly Mg^2+^.^26, 27^ Hence, the stability of single-chain amphiphile membranes towards Mg^2+^ is beneficial for prebiotic compartmentalization scenarios where RNA or RNA-like molecules are hypothesized to have been responsible for the information storage and catalytic functions that today rely on DNA and protein enzymes, respectively.^59, 60^

We compared the stability of coacervate-supported and free vesicle membranes to 25 mM of Mg^2+^ by evaluating their permeability to FITC-Dex 4k. We prepared the experiments by adding the FITC-Dex 4k and Mg^2+^ solution at the same time on the coverslips and let them sit for 24 hours on the bench in the dark for magnesium stability assessment. As a control experiment, we tested the membrane stability 1 day after the addition of solute only (no Mg^2+^). The vesicles and CSM composed of both POPC+OA (50 mol% each) and pure OA membrane didn’t collapse after 1 day but only CSM sample has lower fluorescence intensity of FITC-Dex 4k in general as compared to vesicle membrane (Fig. 4 all panels on the left). In contrast, when 25 mM of Mg^2+^ was added to the samples, both POPC+OA (50 mol% each)- and OA-vesicles formed aggregates instantly (Fig. 4 B and F). On the other hand, the CSMs had better magnesium stability even at 24 hours after the addition of 25 mM Mg^2+^ and FITC-dextran 4k (Fig. 4 D and H).

Clusters/aggregates were observed for both vesicle and CSM samples due to the disruptions from interactions between divalent ions and oleic acid.^61, 62^ We calculated the relative fluorescence intensities 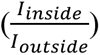 at different time points (1, 12 and 24 hours) among different samples (Fig. 5 A and B). In the absence of magnesium, we observed significantly different relative fluorescence intensities between vesicles and CSM, where vesicles allowed permeation of solute, but CSM didn’t and stayed stable even after 24 hours. In the presence of magnesium, both POPC+OA (50 m% each) and OA-vesicles aggregated and their lumen sizes decreased substantially, such that we couldn’t calculate the relative fluorescence intensities. However, we observed most of the membrane in both POPC+OA (50 m% each) - and OA-CSM samples survived at 24 hours after the addition of 25 mM of Mg^2+^. When increasing the Mg^2+^ concentration to 50 mM, the POPC+OA (50 m% each)-CSM still appeared to have good barrier function, resisting entry of the labeled dextran. On the other hand, OA-CSM started to be influenced more and to show some leakage of solute; hence, the average of relative fluorescence intensities shifted to closer to 1 or even higher than 1. The fact that the average of calculated relative fluorescence intensities POPC+OA (50 m% each)-CSM is much lower than the OA-CSM is consistent with the stabilizing effect of phospholipid in the membrane.

**Figure 5.**
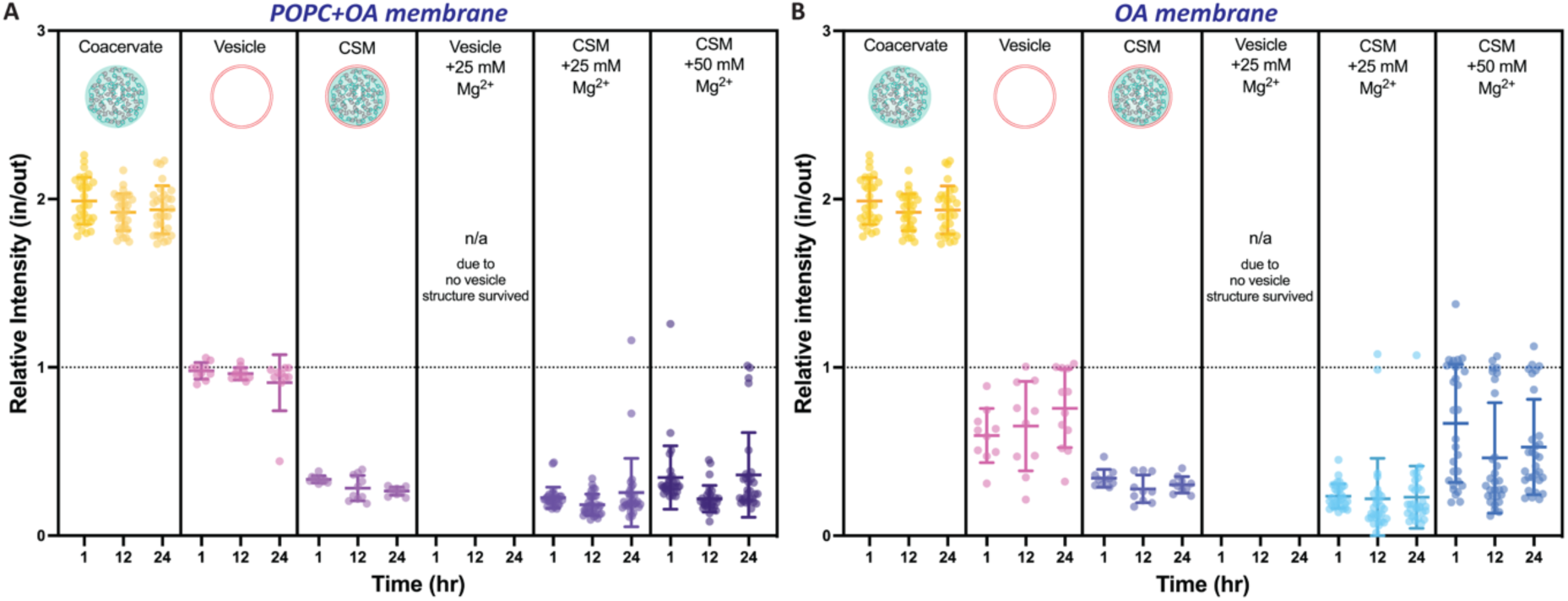
Quantification for coacervate-supported membrane retention of barrier function after treatment with Mg^2+^. Relative fluorescence intensities (in/out) of FITC-Dex 4k by time for various systems, including coacervate, (A) POPC+OA- and (B) OA-vesicles, CSM, vesicle with 25 mM Mg^2+^, CSM with 25 mM Mg^2+^, and CSM with 50 mM Mg^2+^.

### 2.4. Effect of ADP alone on membrane stability to Mg^2+^

Since the anionic component of our coacervates, ADP, binds Mg^2+^ with relatively high affinity^63^, we also tested the protective effect of ADP alone and of the coacervate sample’s dilute supernatant phase. The total [ADP] in our system is 6.67 mM, or 20 mM of negative charge (20 mMc) from ADP, assuming the fully-deprotonated form having -3 charge per molecule at our pH of 8.7. Most of the ADP is sequestered within the coacervate, with only 1.68 mM (5.04 mMc) remaining in the dilute phase. Vesicles formed in charge concentrations of 1.68 mM and 6.67 mM ADP without any polyamine –concentrations that correspond to the measured [ADP] in the dilute phase and the total [ADP]– had good stability to 25 mM Mg^2+^, and excluded the labeled dextran well compared to vesicles in ADP-free solution although not quite as well as coacervate-supported membranes (Supplementary Figure 13). In contrast, vesicles formed in the dilute phase of the coacervate system had become leaky one hour after the addition of 25 mM of Mg^2+^. The difference between dilute phase and 1.68 mM ADP control experiment is that the dilute phase also contains a small amount of the polyamine, which can be expected to compete with Mg^2+^ for binding to the ADP phosphate groups and is more representative of the environment surrounding our coacervate-supported membranes. The inner surface of the coacervate-supported membranes has much higher polyamine concentration that more than compensates for its higher local [ADP]; this is apparent for example in the positive zeta potential for coacervates formed at 1:1 charge ratio (Figure 3C) and is expected based on the difference in translational entropy for the long polyamine vs the much smaller ADP molecules.^9^ Therefore, while some membrane stabilization may arise due to the Mg^2+^-chelating ability of ADP, we interpret the superior Mg^2+^ stability for fatty acid membranes in contact with coacervate phase as primarily due to membrane stabilization by ion-pairing interactions between fatty acid headgroups and coacervate amine groups (from PAH), as suggested by the reduced lateral fluidity of these coacervate-contacting membranes.

**Figure 6.**
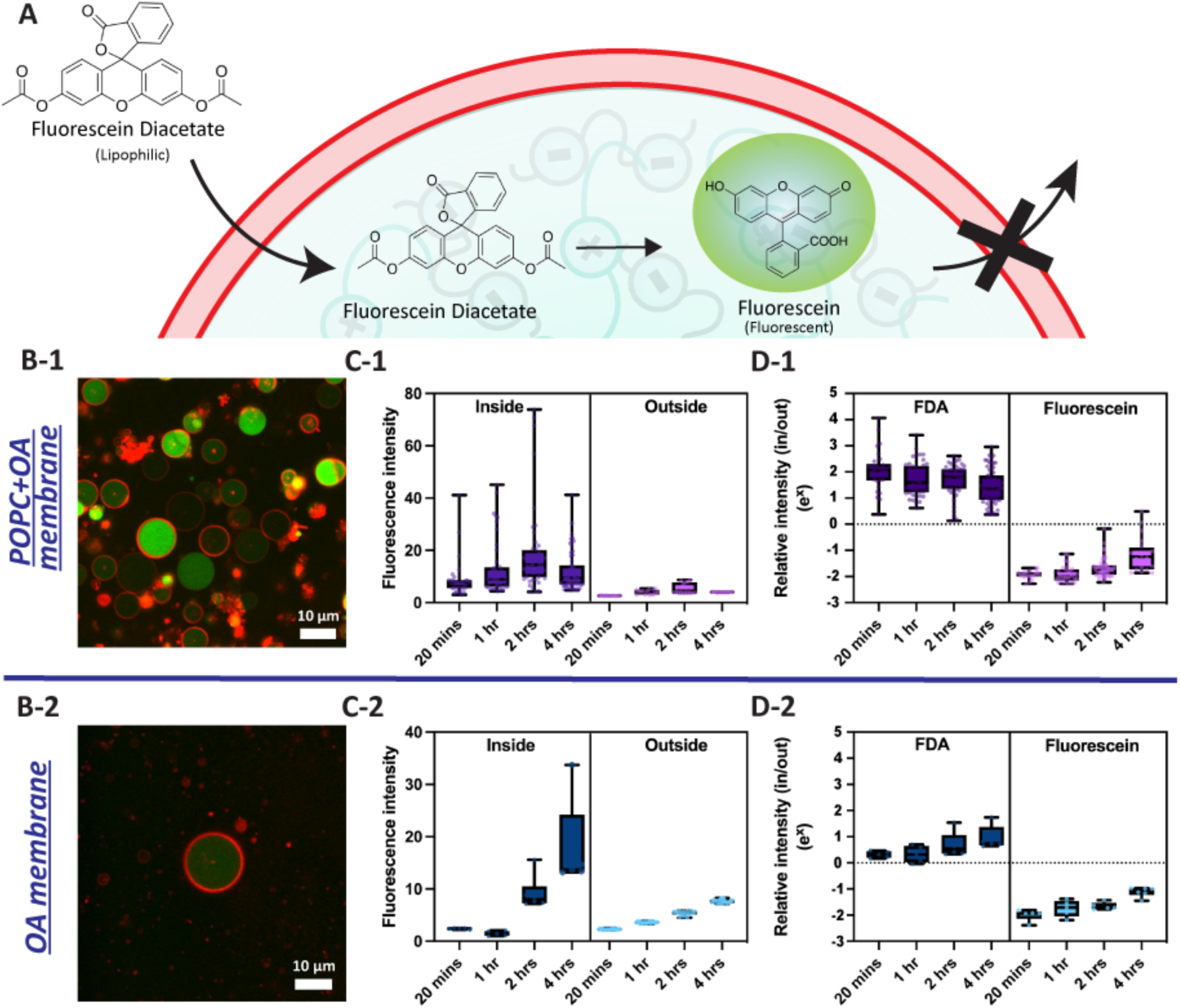
Substrate entry, ester hydrolysis and fluorescent product retention in coacervate-supported membrane hybrid protocells. (A) Non-fluorescent lipophilic fluorescein diacetate (FDA) added externally enters coacervate interiors by diffusion across membranes and is subsequently hydrolyzed to the anionic fluorescent product, fluorescein. Confocal fluorescence images of FDA hydrolysis in POPC+OA- and OA-CSM (B-1 and -2, respectively) 4 hours after addition of FDA. Fluorescense intensities of product (fluorescein) changes over time inside and outside the POPC+OA- and OA-coacervate-supported membranes (C-1 and -2). Comparison of relative fluorescence intensity (in/out) after adding either the nonfluorescent substrate FDA, which generates the fluorescent product upon ester hydrolysis (left side), or the fluorescent product fluorescein (D-1 and D-2).

### 2.5. Substrate entry, ester hydrolysis and fluorescent product retention in coacervate-supported membrane hybrid protocells

Intracellularly, non-specific esterases and some free amino acids are able to cleave ester bonds, a capability that is often leveraged to deliver small molecules into biological cells for imaging by dosing the cells with an ester-protected version of the molecule. After this molecule permeates the cell membrane, hydrolysis converts the neutral ester moieties to negatively-charged carboxylates. The anionic product is unable to permeate the membrane and becomes concentrated inside. Since this process requires both intracellular esterase activity and an intact membrane, it is commonly used to determine whether cells are alive or dead. Our hybrid protocells consisting of polyamine/ADP-rich coacervate “cytoplasm” and POPC+OA or pure OA membranes are able to pass this primitive cell viability test due to the highly concentrated amine moieties in the coacervate interior and the strong barrier property of the surrounding membranes, which allow the uncharged acetate-protected small molecule FDA to pass, while retaining the anionic product (fluorescein)^64^, which is fluorescent and can be observed under fluorescence microscopy (Fig. 6A, B). We note that other related molecules are also used for this purpose, and are preferred for lower (physiological) pH and when slower leakage of the product from live cells is needed.^65^

After external addition of the nonfluorescent substrate FDA, coacervate cores coated with either POPC+OA- or OA-CSMs showed increased fluorescence intensity from interior fluorescein accumulation over time (Fig. B-1 and -2, Supplementary Figure 14). Variability in product generation (green fluorescence intensity) between structures within a sample, evident in Figure 6A, appears to be correlated with the concentration of coacervate interiors, which varies some within the population due to membrane assembly outpacing coacervate equilibration during CSM formation. Transmitted light images show the most-active structures are also those with higher refractive index contrast with the surrounding medium; their higher polyamine content is likely responsible for their more rapid product generation (Supplementary Figure 15). Such variability is potentially of interest for future investigations where differences in reactivity could impact “fitness” under selection pressure. The timescale of internal fluorescein accumulation was much more rapid for POPC+OA than OA-only CSMs (compare Figure 6C-1 and C-2). We interpret this as a consequence of the stronger barrier function of these OA-only CSMs, as observed for the fluorescent solutes in Figure 2. Fluorescence intensity also increased somewhat outside of the membrane-coated coacervates. Control experiments indicate that not only coacervate droplets (with or without buffer), but also tris buffer alone, can catalyze the formation of fluorescein from FDA (Supplementary Figure 16). This effect of tris buffer has been reported previously by Wanandy et. al.^66^ and Clarke et. al.^67^ We did not try to remove tris buffer for CSM due to the reported strong pH-sensitivity of oleic acid vesicle formation.^68^ Despite some hydrolysis occurring in the dilute phase due to tris, the fluorescein intensity was much higher in the coacervate interiors. Since the CSMs do not allow fluorescein to enter at short times (Figure 2, top row), and only modest leakage is found at long times (Figure 6D and Supplementary Figure 17), this effect was not due to simple partitioning of fluorescein formed externally and is due to the reaction occurring inside. Direct comparisons of the intensity ratios after addition of FDA or fluorescein make clear that fluorescein accumulates more quickly inside protocell interiors when added as the membrane-permeable, nonfluorescent FDA than as the anionic, less membrane-permeable fluorescein (Figure 6D). Although obviously a far stretch from living biological cells, these CSM-based hybrid protocells were to mimic their behavior in this crude cell viability assay through the combination of reactive coacervate cores and solute-gating boundary membranes.

## 3. Conclusion

Fatty acid and fatty acid/phospholipid blended membranes templated by coacervate droplets were self-assembled via a modified gentle hydration approach in which preformed coacervate droplets were used as the hydrating liquid for dried lipid films. The resulting coacervate-supported membranes provided an effective boundary membrane for the droplets, and showed barrier function as good as or better than traditional vesicles formed from the same amphiphiles. This observation differs from previous work from our lab^48^ and others^47^, which reported greater permeability for coacervate-supported membranes. Differences in coacervate and lipid compositions, and/or in the lipid assembly process are presumably responsible for the differences in membrane permeability to solutes observed between these publications. The relatively wide range of membrane permeability properties observed across these different systems promises the possibility of design-based control over permeability once the factors controlling it are better understood. Important factors include membrane composition, coacervate composition, and the way amphiphiles and coacervates come into contact (e.g., coacervates present during lipid hydration, soluble lipids added, etc). For the PAH/ADP coacervate, POPC/OA or OA membrane system studied here, attractive interactions between negatively-charged OA headgroups and the positively-charged amine sidechains of PAH in the coacervate droplets were important in formation and properties of the coacervate-supported membranes. Membrane stabilization through these charge-charge interactions with the coacervate interior was apparent in the decreased lipid diffusion and increased barrier function of the coacervate-supported membranes, particularly in the resistance to disruption by externally added Mg^2+^.

Greatly improved Mg^2+^ stability afforded CSMs by interactions with the coacervate core could be beneficial for an RNA World type scenario where Mg^2+^ is needed for proper folding and activity of functional RNAs.^17, 18, 27^ A somewhat unexpected possible challenge for the hybrid protocells described here is that their membranes’ barrier function may be too effective, potentially hindering the entry (and egress) of necessary small molecule “fuel” (and waste). We note that the solutes studied here were either polyanionic (fluorescein, calcein, U15 RNA) or relatively large (dextran 4k), which enabled us to compare membranes with and without coacervate cores; passage of relatively smaller, less charged solutes is less inhibited. We demonstrated this using the lipophilic substrate, FDA, and which readily penetrated the coacervate-supported membranes, was reacted to form the anionic fluorescent product, and was subsequently trapped within the hybrid protocells. Such processes begin to mimic living cells and could be considered as a very primitive step towards abiotic metabolism in the hybrid membrane-coacervate protocells. It will be interesting in future work to explore whether coacervates can stabilize membrane assembly from shorter, more soluble lipids that may be unable to form functional membranes on their own, as such molecules may have been more available prebiotically, and could potentially provide membranes with greater permeability than the OA and POPC+OA used here.

## 4. Materials and Methods

### 4.1. Materials

Poly(allylamine hydrochloride) (PAH, MW 17.5 kDa), Adenosine-5’-diphosphate (ADP), MgCl_2_, Tris (Trizma Base), Fluorescein, Calcein, Nile red, Propidium iodide, FITC-Dextran (MW 4 kDa), and Cy5 labeled U15 (MW 5 kDa), fluorescein diacetate were purchased from Sigma-Aldrich. 1-palmitoyl-2-oleoyl-sn-glycero-3-phosphocholine (16:0-18:1 PC, POPC), and 1,2-dioleoyl-sn-glycero-3-phosphoethanolamine-N-(lissamine rhodamine B sulfonyl) (ammonium salt) (18:1 Liss Rhod PE, Rhod-PE) were purchased from Avanti polar lipids. Oleic acid was purchased from Nu Chek Prep. PAH and ADP stock solutions were prepared in 5 wt% and 100 mM respectively. Both solutions were pH corrected to around pH 7.4 using 1 M NaOH. 1 M Tris buffer was pH corrected to 8.7 with 1 M HCl. All lipids were aliquoted and stored at -80℃, except oleic acid at -20℃.

### 4.2. Preparation of Coacervate-Supported Membranes

First, a lipid film was prepared as follows. Lipid chloroform solution with the desired composition of 2.5 mg/mL fatty acid and/or phospholipid was prepared in glass test tube and dried with argon gas to form a thin lipid film on the glass. The lipid films were then kept under vacuum for 6 hrs in order to make sure that all the chloroform was removed. Meanwhile, a coacervate sample was prepared separately, to be used to hydrate the lipid film. Coacervate samples were prepared by adding reagents in the order: HPLC grade water, Tris buffer (stock 1 M, final concentration 250 mM, pH 8.7), ADP (6.67 mM final concentration, which corresponds to 20 mM of negative charge from ADP, based on assumption of -3 charge per molecule at our pH), and PAH (20 mM charge concentration) for 1 to 1 charge ratio coacervates. We use units of mMc to indicate mM in charge from a particular molecule, for ease of clarifying whether components of complex coacervates are charge matched). A turbid solution was obtained immediately after adding PAH, indicating phase separation to form the coacervate droplets. After pipette mixing 10 times, the coacervate suspension was added to the dried lipid film. Then, the sample was heated at 45 °∁ for 24 hrs for lipid assembly. For comparison, samples of vesicles without coacervates were prepared with 250 mM Tris buffer (pH 8.7) in place of the coacervate suspension and otherwise treated the same as the samples with coacervates.

### 4.3. Surface charge measurements

Zeta potential measurements were conducted using a Malvern DLS-Zetasizer. Coacervates were diluted by 4 times and freshly prepared right before the measurements. Each sample was measured three times at 25 °∁. Vesicle and coacervate-supported vesicles samples were diluted by 4 times and then measured three times at 25 °∁.

### 4.4. Fluorescence Recovery after Photobleaching

FRAP studies were carried out using a confocal fluorescence microscope (Leica TCS SP5) equipped with an HCX PL APO CS ×63.0/1.40 NA oil UV objective. FRAP experiments were conducted on membranes of interest that contained 0.05 mole% rhodamine-labeled lipids using excitation at 543 nm. For samples without coacervates (vesicle and vesicle prepared with dilute phase), we performed bi-directional scanning to catch the fast recovery from photobleaching. A 2 µm circular area of interest in the membrane (moving to the top of the vesicle in order to obtain a homogeneous area) was bleached using lasers (458, 476, 488, 514, 543, and 633 lasers) with all 100 % power for 10 frames and followed by 200 post-bleach frames. For vesicle (without coacervate) samples, bidirectional scanning was used in order to catch the faster recovery of lipid membrane in vesicles.

Background noise was corrected by measurement of fluorescence intensity in ROIs when all lasers were all turned off, but the respective photomultiplier tube (PMT) was on. A reference ROI was obtained meanwhile to exclude the effect of photobleaching from imaging. Apparent diffusion coefficient was calculated as in a previous paper that discusses photobleaching lipid membrane around coacervate^1^.

Fluorescence intensity (F_N_(t)) was normalized using the double normalized method (**Equation 1**)^.2, 3^

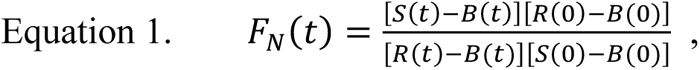

where S(t) is the average fluorescence intensity of the ROI chosen for analysis, R(t) is the average florescence intensity of the reference ROI, and B(t) is the average background fluorescence intensity.

Date was fit using the single exponential recovery function (**Equation 2**) in Igor Pro 8.

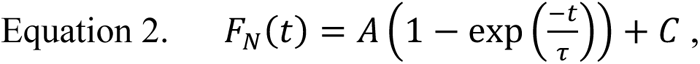

where A is the mobile fraction of the fluorescence probe that id able to recover, C represents the y-intercept of the recovery curve, and τ represents the fluorescence recovery time constant. The recovery halftime (τ_1/2_) was the calculated using **Equation 3**.

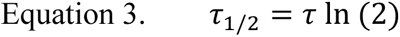

Last, the apparent diffusion coefficient (D_app_) was calculated using **Equation 4**.

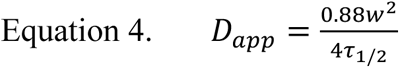

At least 10 FRAP experiments were done for each trial (i.e. 10 droplets bleached) and three independently-prepared trials were done for each sample.

#### Permeability Examination of CSMs, coacervates and liposomes

Permeability studies were performed using the same confocal microscope (Leica TCS SP5) and HCX PL APO CS ×63.0/1.40 NA oil UV objective. To conduct the permeability study, 20 µL of CSMs or coacervates solution were added to the coverslips and then 1 µL of desired solute was added right after to achieve final concentration of 5 µM for fluorescein, calcein and FITC-Dex 4k, 10 µM for Cy5-U15 RNA, and 20 µM for nile red and propidium iodide. Among those solutes, Cy5-U15 has much stronger accumulation into the coacervates, and due to its interactions with the coacervate components Cy5-U15 takes longer equilibrate across the population of droplets. Therefore, we waited 1 day as compared to other samples only waited for 20 minutes. At least five images were taken 20 minutes (for fluorescein, calcein, nile red, propidium iodide, and FITC-dextran 4kDa) or 1 day (for Cy5-U15 RNA) after the addition of solute to coacervate samples using confocal microscopy. Around twenty to fifty images were taken for the vesicle and CSM samples. Background of signal was obtained with the same imaging settings from the samples without solute addition. ImageJ (Fiji) was used to obtain the fluorescence intensities inside and outside (average of five ROIs from 4 corners and middle of the image). Average intensity ratio was calculated by subtracting the background intensity when no fluorescent solute added. All experiments were done in triplicate.

### 4.5. PAH and ADP concentration in dilute phase

After the preparation of coacervate solution as above, centrifuged the solution at 9.3 rcf at 20°∁ for 10 mins using Eppendorf centrifuge model 5415R. Took out the dilute phase gently and measure the absorbance of ADP at 260 nm using Agilent model 8453 UV-Vis spectrometer. Calibration curve was performed to confirm the exact concentration of ADP concentration in dilute phase. PAH concentration in dilute phase was examined using Alexa 488 labeled PAH. Same amount of labeled PAH was added to all samples and characterized using Jobin Yvon Fluorimeter model FL3-21 excited using 488 nm and collected the emission between 500-600 nm. Percentage of PAH left in dilute phase were calculated by *Fluorscence intensity of dilute phase/Fluorescence intensity of bulk* at emission at 515 nm (maximum of emission).

### 4.6. Magnesium stability test of membrane

After gentle hydration, instead of agitating the sample immediately, 250 µL of solution (mainly dilute phase) was removed from 1 mL sample in the test tube. Agitated the sample and took 22.5 µL to the coverslips following with 7.5 µL 100 mM MgCl_2_ prepared in coacervate dilute phase, and 1.5 µL of 100 µM FITC-Dex 4k. The samples were then imaged by confocal fluorescence microscopy after 1, 12, and 24 hours after the preparation to determine uptake of the labeled dextran into the vesicle lumen/coacervate interiors. All experiments were done in triplicate. We noticed the osmolarity change when adding the stock MgCl_2_ to the lipid samples and caused the flocculation among vesicles. Therefore, we minimized the osmolarity change by adding the MgCl_2_ prepared in dilute phase of coacervate which minimize the osmolarity shock that may cause to the membrane (Supplementary Figure 12). In addition, to exclude the factor for concentration gradient within sample caused by the addition of solute and magnesium on coverslip, we also performed the magnesium stability test in Eppendorf tube as well and observed no significant difference in average relative fluorescence intensities between two methods (Supplementary Figure 13). However, more clumping was observed when the stability test was done in the tube.

### 4.7. FDA hydrolysis

After the preparation of CSM or coacervate solutions, we added 50 µL of the solution on the coverslip and then followed with 1 µL of 1 mg/mL FDA solution prepared in acetone (final FDA concentration is ∼47.1 µM). We took images every minute for 20 minutes, as well as time points at 1, 2, and 4 hours. Around ten images were taken for image analysis.

Background of signal was obtained with the same imaging settings from the samples without solute addition. ImageJ (Fiji) was used to obtain the fluorescence intensities inside and outside (average of five ROIs from 4 corners and middle of the image). Average intensity ratio was calculated by subtracting the background intensity when no fluorescent solute added. All experiments were done in triplicate. As the control experiment for fluorescein leakage. We performed the fluorescein permeability test with higher concentration (higher than the concentration we usually use for permeability test) to match with the FDA concentration (assume all FDA hydrolyzed into fluorescein), which was 47.1 µM. Images were collected and analyzed in the same way.

## Supporting information

Suppenmentary Information

## Acknowledgement

This work was supported by the National Science Foundation Grant 1844313—RoL: RAISE: DESYN-C3 and Huck Institutes of the Life Sciences at Penn State University through the Huck Innovative and Transformational Seed Grant (HITS). Content is the responsibility of the authors and does not represent the views of the NSF or Huck Institutes. The authors thank Kate Adamala for helpful discussions about oleic acid membranes.

