## Supplementary material for "Hybrid Protocells based on Coacervate-Templated Fatty Acid Vesicles combine Improved Membrane Stability with Functional Interior Protocytoplasm": Suppenmentary Information

### Supplementary Figures and Tables.

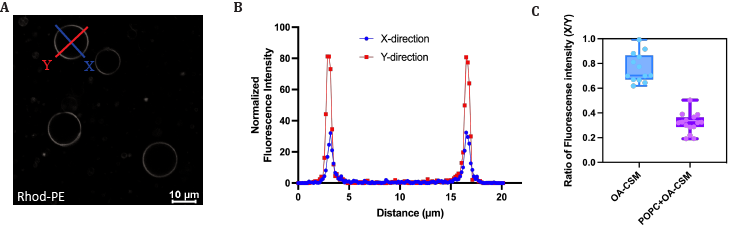

Figure S1**. Fluorescence polarization for blended phospholipid/fatty acid coacervate-supported membranes (POPC+OA-CSM).** (A) Confocal fluorescent microscopic images of POPC+OA-CSM (0.05 mol% Rhod-PE). Difference of fluorescence intensity between x- and y- direction were observed when the polarizers are inserted in confocal microscopy. (B) Example of normalized fluorescence intensity of x-/y-directions. (C) Statistical data for polarization ratio out of 12 OA-CSMs and 14 POPC+OA-CSMs.

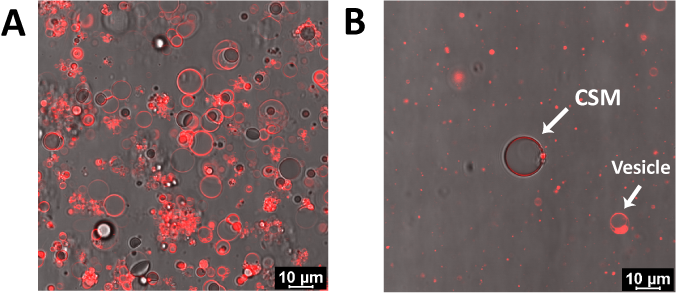

Figure S2**. Example of microscope images for (A) fatty acid-blended (POPC+OA-CSM) and (B) fatty acid-only (OA-CSM).** Transmitted light ^6^ and rhodamine-PE lipid confocal fluorescence channels are overlaid. The yield of OA-CSM was relatively lower than POPC+OA-CSM determined by the average amount of structure within one frame.

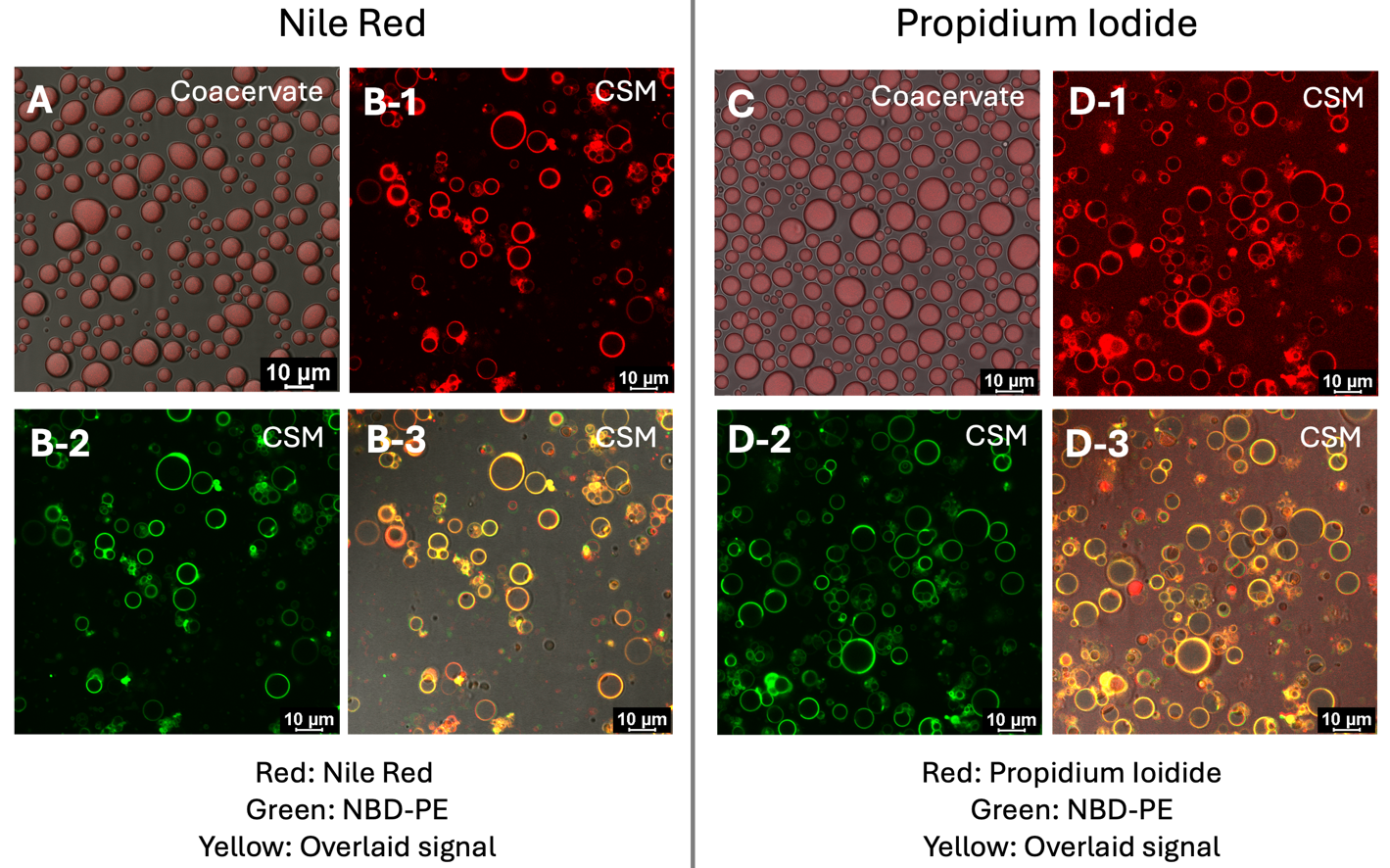

Figure S3**. Addition of lipophilic (Nile Red) and cationic (Propidium iodide) solute to PAH+ADP coacervate and POPC+OA-CSMs after 20 minutes.** Both solutes (A and C) partition within the coacervate droplets when there is no membrane around. On the other hand, both solutes show interactions with the membrane. While Nile Red mainly locates within the membrane, Propidium iodide shows fluorescence intensity at both bulk phase and around membrane. Nile red prefers to get in the membrane due to its lipophilic property. Cationic propidium has strong interaction to membrane due to the electrostatic interaction since the oleic acid in membrane is negatively-charged at the pH we used.

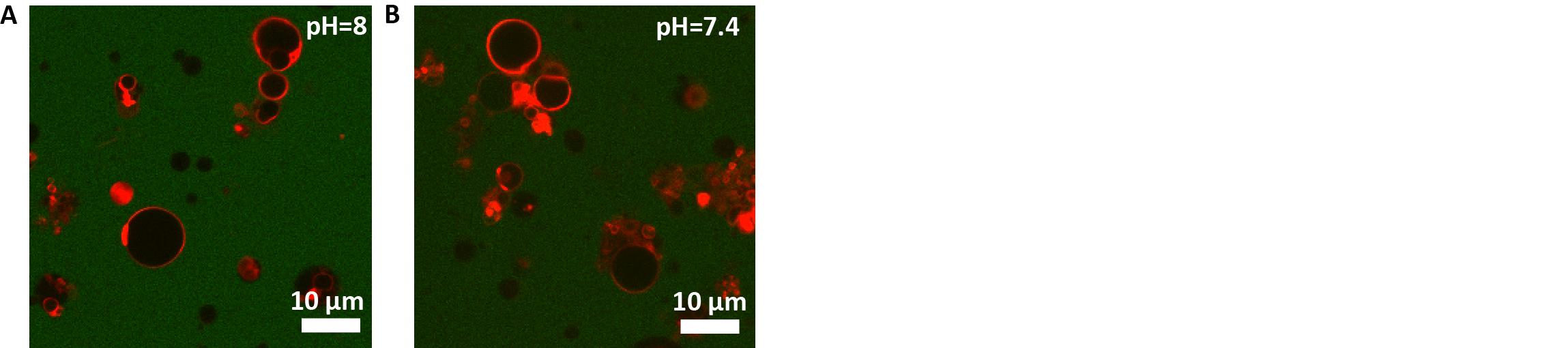

Figure S4**. POPC+OA-CSMs prepared at pH 8 and 7.4 are impermeable to calcein at 20 minutes** Permeability test with calcein for POPC+OA-CSMs prepared at pH 8 and 7.4, which are relatively lower pH for oleic acid and physiological pH. Both CSMs showed to be able to prevent the entry of calcein after 20 mins.

| Sample 1 | Sample 2 | p-value |
| --- | --- | --- |
| **Fluorescein** |  |  |
| POPC+OA-vesicle | POPC+OA-CSM | < 0.0001, **** |
| OA-vesicle | OA-CSM | < 0.0001, **** |
| POPC+OA-vesicle | OA-vesicle | 0.0005, *** |
| POPC+OA-CSM | OA-CSM | 0.7958, ns |
| **Calcein** |  |  |
| POPC+OA-vesicle | POPC+OA-CSM | < 0.0001, **** |
| OA-vesicle | OA-CSM | < 0.0001, **** |
| POPC+OA-vesicle | OA-vesicle | < 0.0001, **** |
| POPC+OA-CSM | OA-CSM | < 0.0001, **** |
| **FITC-Dex 4k** |  |  |
| POPC+OA-vesicle | POPC+OA-CSM | < 0.0001, **** |
| OA-vesicle | OA-CSM | < 0.0001, **** |
| POPC+OA-vesicle | OA-vesicle | < 0.0001, **** |
| POPC+OA-CSM | OA-CSM | 0.0950, ns |
| **Cy5-U15** |  |  |
| POPC+OA-vesicle | POPC+OA-CSM | < 0.0001, **** |
| OA-vesicle | OA-CSM | < 0.0001, **** |
| POPC+OA-vesicle | OA-vesicle | < 0.0001, **** |
| POPC+OA-CSM | OA-CSM | 0.1512, ns |

Table S1**. Two tailed p-values of average relative intensity ratio for vesicle and CSM with two lipid compositions (POPC+OA and OA) tested with four solutes.**

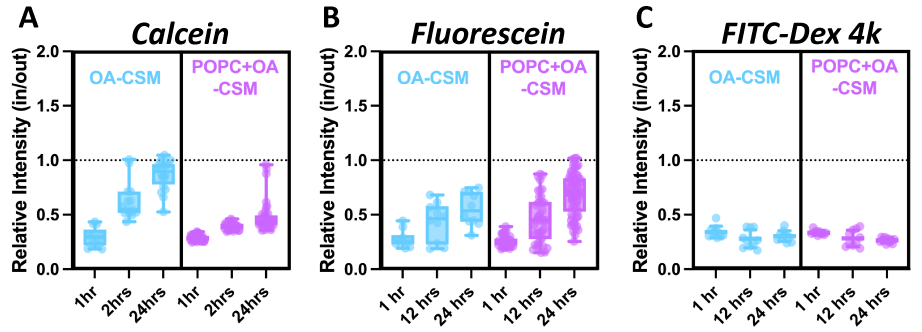

Figure S5**. Relative intensity ratios (in/out) of OA- and POPC+OA-CSMs at 1, 12, and 24 hrs after the addition of (A) calcein (anionic), (B) fluorescein (anionic), and (C) FITC-Dex 4k (neutral).**

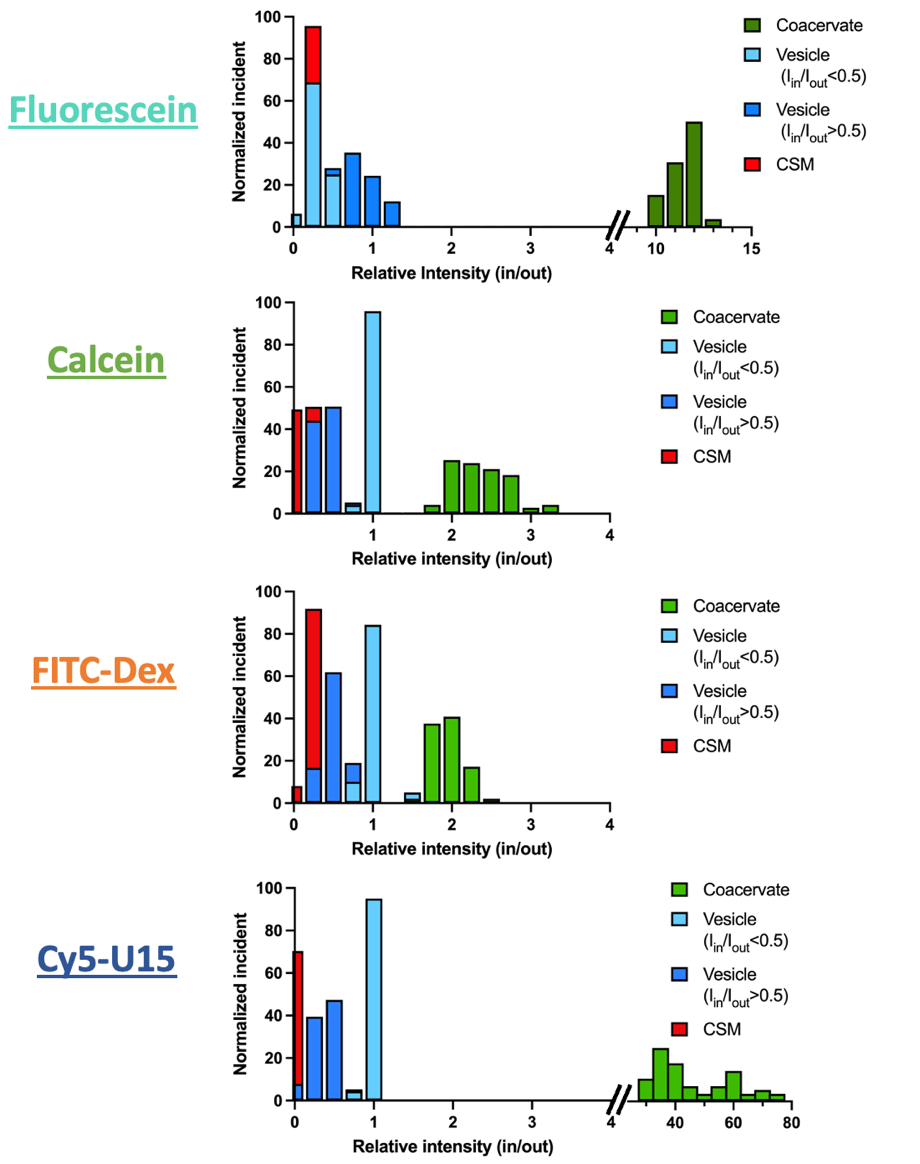

Figure S6**. Relative intensity (in/out) values distribution of coacervate, POPC+OA-vesicle, separating into penetrated (relative intensity > 0.5) and non-penetrated (relative intensity < 0.5), and POPC+OA-CSM.**

|  | **POPC+OA-Coacervate-supported membrane (CSM)** | | | **Vesicles** | | |
| --- | --- | --- | --- | --- | --- | --- |
|  | # impermeable | # permeable | Percent of structures permeable to solute | # impermeable | # permeable | Percent of structures permeable to solute |
| Fluorescein | 136 | 2 | 1.4% | 48 | 82 | 63.1% |
| Calcein | 266 | 1 | 0.4% | 92 | 260 | 73.9% |
| FITC-Dex 4k | 211 | 0 | 0% | 57 | 317 | 84.8% |
| Cy5-U15 | 153 | 0 | 0% | 51 | 257 | 83.4% |

Table S2**. Statistical data for solute permeability for POPC+OA-CSM and vesicles (without coacervates).** Percent permeable as determined by relative solute concentration with a cutoff threshold of 0.5. Relative solute concentration (Intensity inside/intensity outside) higher than 0.5 was considered permeable, and lower than 0.5 is considered impermeable.

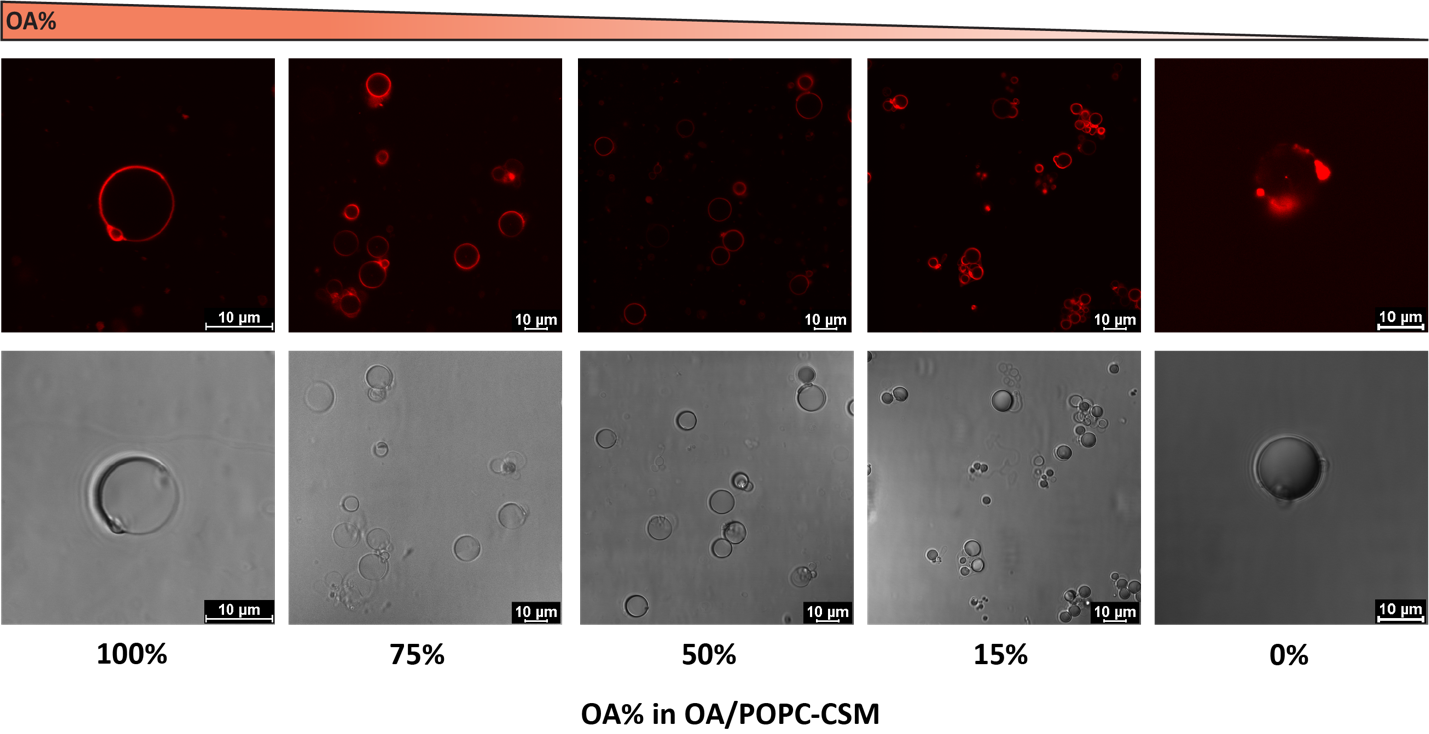

Figure S7**. Effect of varying the OA% in OA/POPC-membrane on formation of CSMs.** For CSMs composed of OA% from 100-15% all had complete membrane structures for further FRAP experiments. However, pure POPC composition resulted in lipid aggregates at PAD+ADP coacervate droplets.

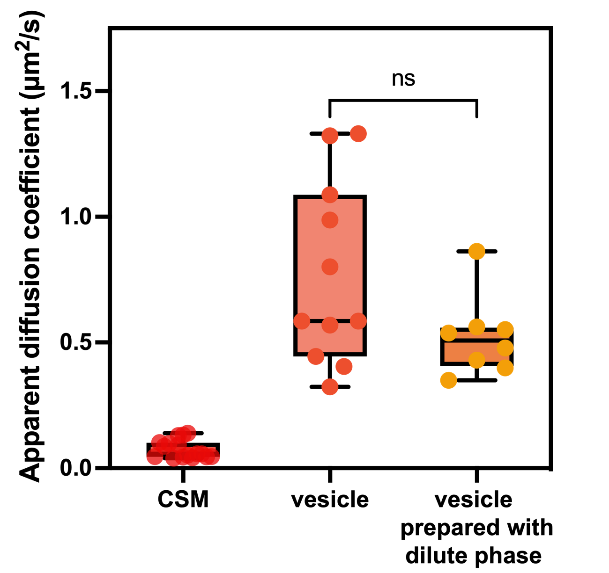

Figure S8**. Comparison of lipid mobility in POPC+OA (50% OA) -CSM, -vesicles, and -vesicles prepared in dilute phase of the coacervate samples.** Apparent diffusion coefficients for labeled lipid molecules in these membranes were determined by fluorescence recovery after photobleaching (FRAP). Apparent diffusion coefficients in vesicles formed in Tris buffer (250 mM at pH 8.7) or in dilute phase of the coacervate samples were not significantly different, indicating that the much smaller apparent diffusion coefficient for lipids in CSM is mainly caused by interactions with the coacervate interior rather than ADP or PAH molecules present in the dilute phase.

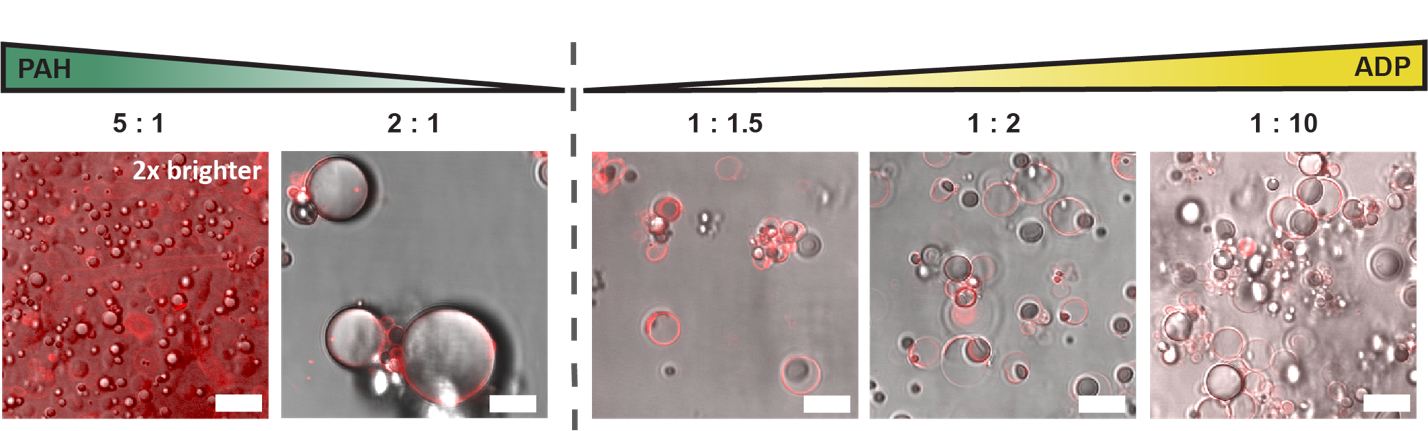

Figure S9**. POPC+OA (50 mol% for each) lipid assembly at coacervates with various PAH:ADP charge ratios.** The relative amount of PAH and ADP were varied from fivefold excess PAH to tenfold excess ADP (charge ratios of PAH:ADP are given in the figure) Scale bar 10 μm.

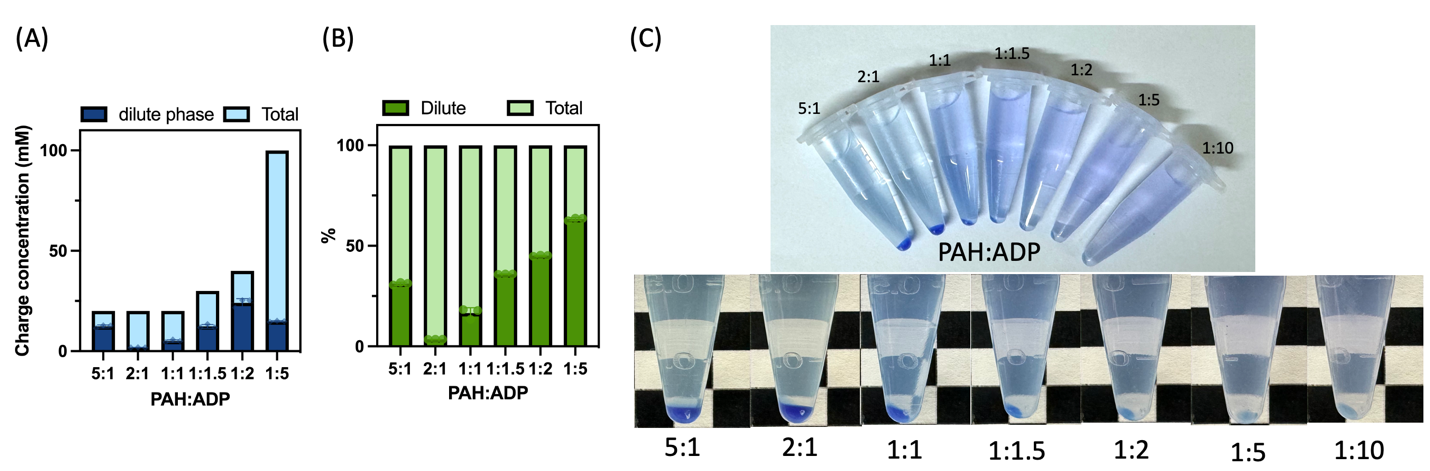

Figure S10**. Impact of PAH:ADP charge ratio on relative phase volumes and coacervate component partitioning.** Measurements of ADP (A) and PAH (B) in dilute phase. 1:1 charge ratio was prepared with each 20 mM charge concentrations of ADP and PAH. Coacervate volume with different charge ratios (C). Varying the charge concentration of polyelectrolytes changes the amounts of polymers left in dilute phase and the volume of coacervate phases (bromophenol blue dye was added to aid visualization).

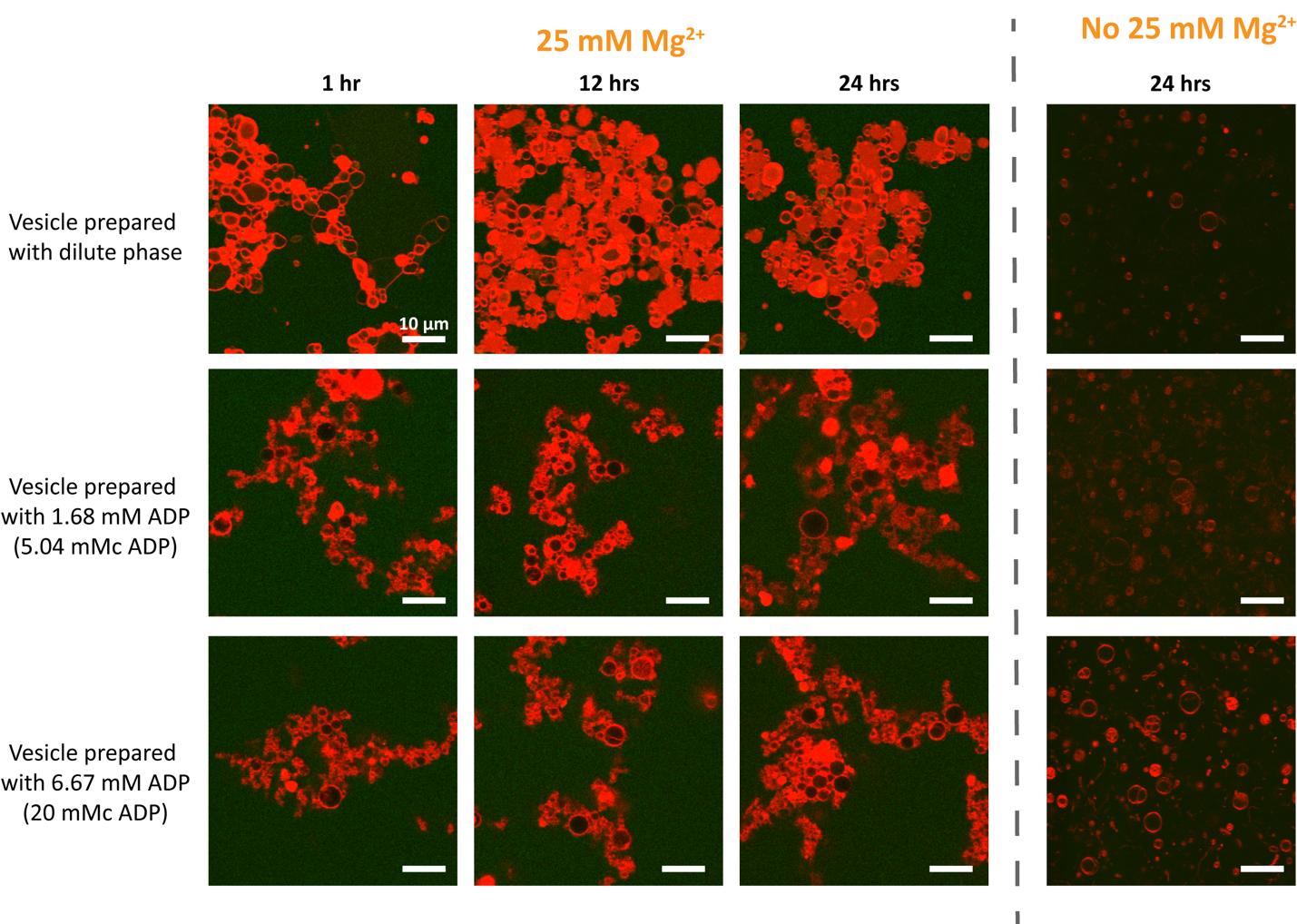
Figure S11**. 25 mM Mg^2+^ stability test of POPC+OA-vesicles prepared with various conditions.** In order to understand the impact of molecules left in dilute phase influencing the membrane stability, we performed the magnesium stability test with vesicles in dilute phase, and in controls that lacked the polycationic species (PAH) so could not form coacervates but had either 1.68 mM ADP (5.04 mMc from ADP, corresponding to its concentration in dilute phase) and 6.67 mM ADP (20 mMc from ADP, corresponding to the total amount added when forming coacervates, most of which becomes localized within the coacervate phase) Scale bar 10 μM.

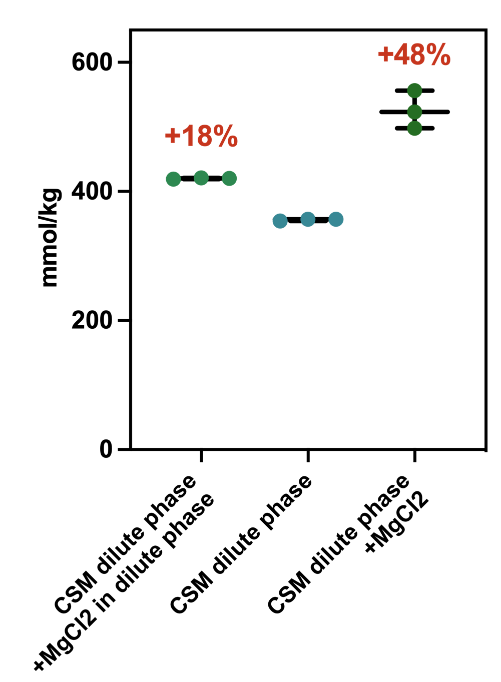

Figure S12**. Osmolarity measurement of the addition of MgCl_2_ stock prepared with just water or with dilute phase of coacervate.** In order to minimize the effect of osmolarity shock, which caused by the imbalance of osmolarities inside and outside the membrane, we performed the osmolarity measurement. Data show the less osmolarity change when we prepared the MgCl_2_ stock solution in coacervate dilute phase as compared to MgCl_2_ stock prepared with HPLC water.

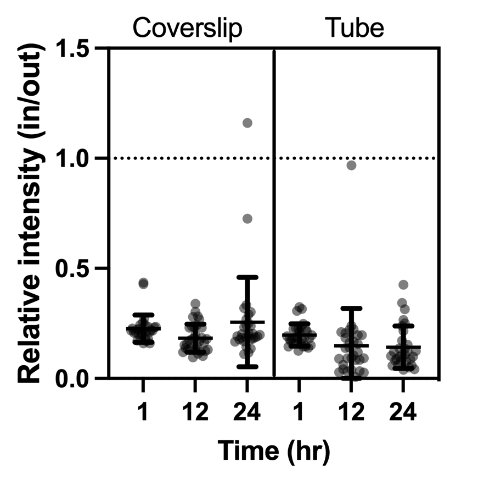

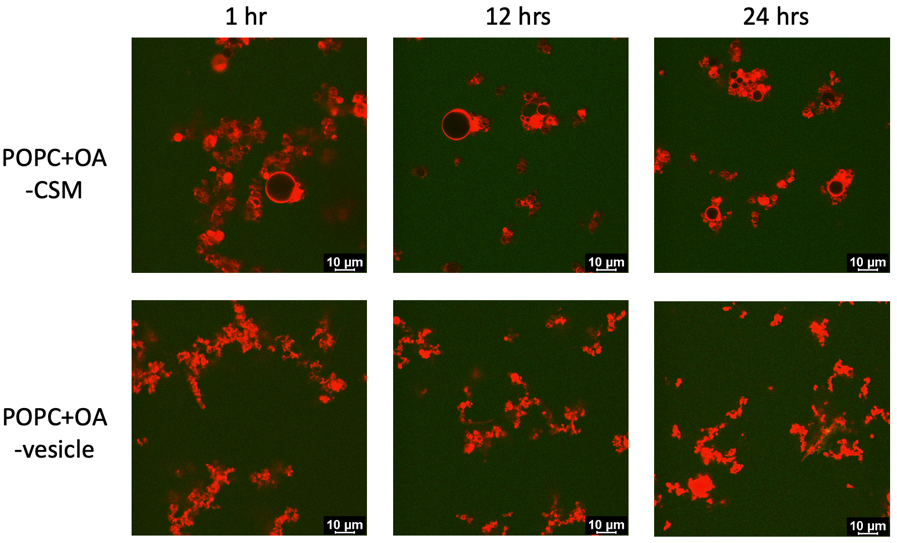

Figure S13**. Confocal microscopic image of 25 mM Mg^2+^ stability test of POPC+OA-CSM and vesicle performed in Eppendorf tube and the comparison of relative intensity (in/out) of POPC+OA-CSM on coverslip and in tube.** To further exclude the impact of concentration gradient due to performing the test on coverslip, we did the magnesium stability test in the tube instead and took out the solution from the tube at different time points (1, 12, 24 hrs). No obvious difference was observed as compared to do the magnesium stability test on the coverslips. POPC+OA-CSM has better magnesium stability compared to vesicles determined by both microscopic images and permeability examination (fluorescence intensity inside and outside).

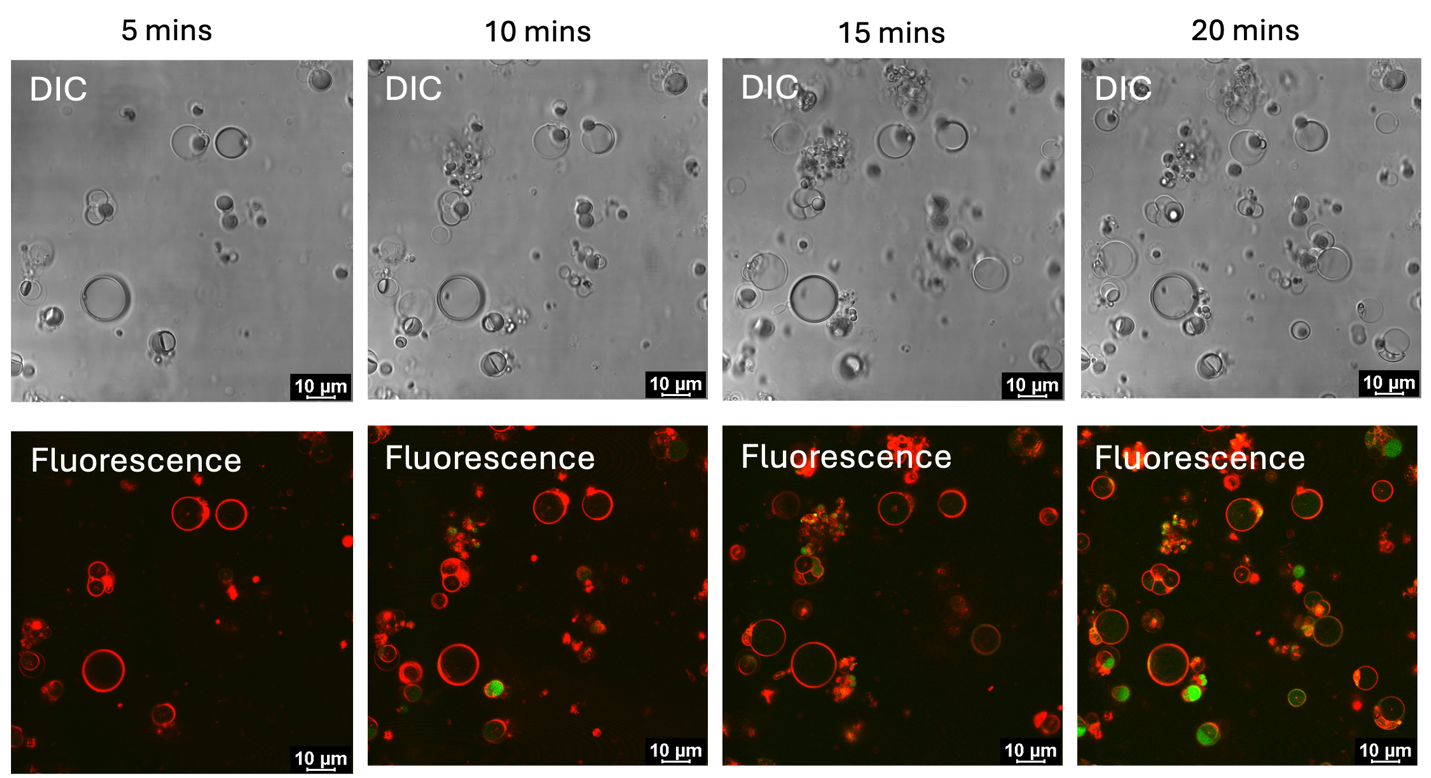
 Figure S14**. FDA added to POPC+OA-CSM.** FDA with final concentration 47.1 µM was added to the POPC+OA-CSM. The product (fluorescein) of hydrolysis accumulated within the membrane since lipophilic FDA simple diffused into the CSMs and was hydrolyzed within the protocytoplasm.

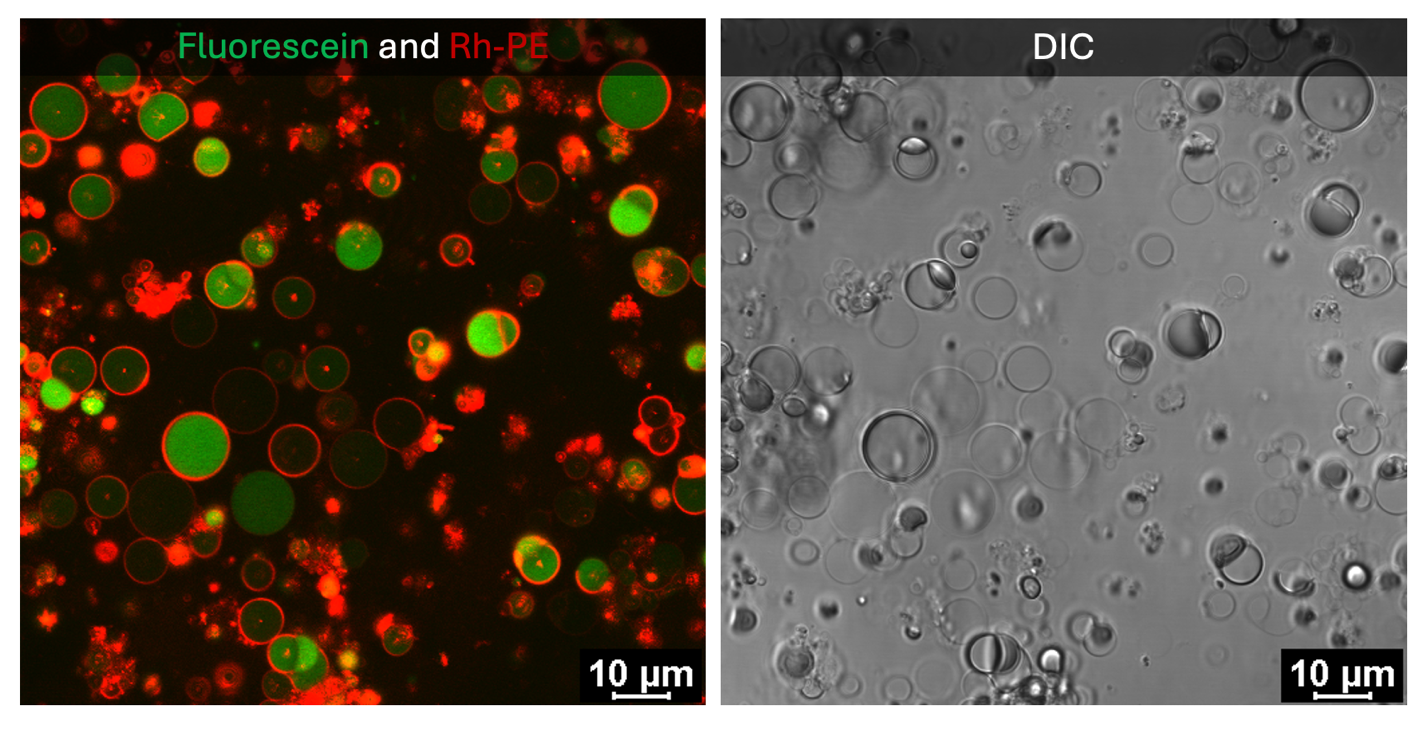

Figure S15**. Confocal microscopic fluorescence and DIC images of FDA added to POPC+OA-CSMs.** We observed that there was heterogeneity of refractive index among CSMs in DIC channel. The refractive index seemed to correlate with the fluorescein intensity. The CSM had higher refractive index also had higher fluorescein intensity.

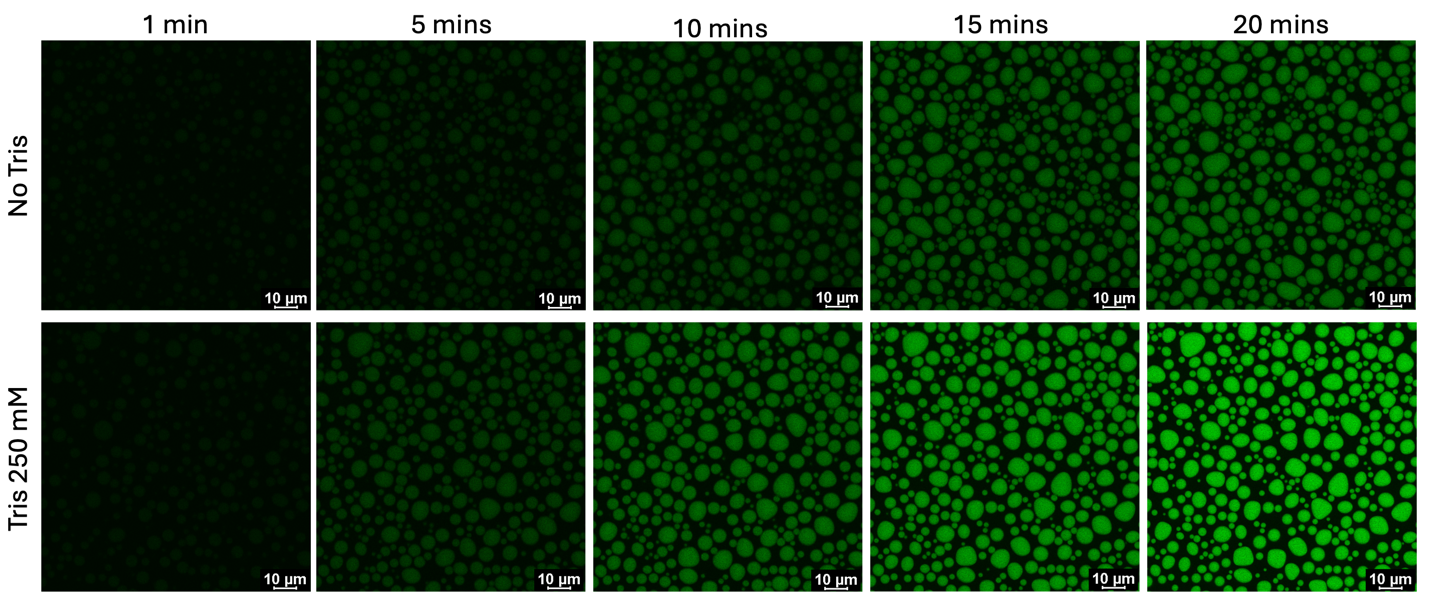
Figure S16**. Fluorescein intensity increase after adding FDA to PAH+ADP coacervates prepared with and without 250 mM Tris buffer.** To understand the effect of tris buffer on the FDA hydrolysis, we added FDA to coacervates prepared with and without the buffer. Although the coacervates prepared with tris buffer had faster FDA hydrolysis, the coacervate prepared without tris buffer also showed substantial increase of fluorescein intensity over time, indicating that the coacervate itself also facilitate the hydrolysis.

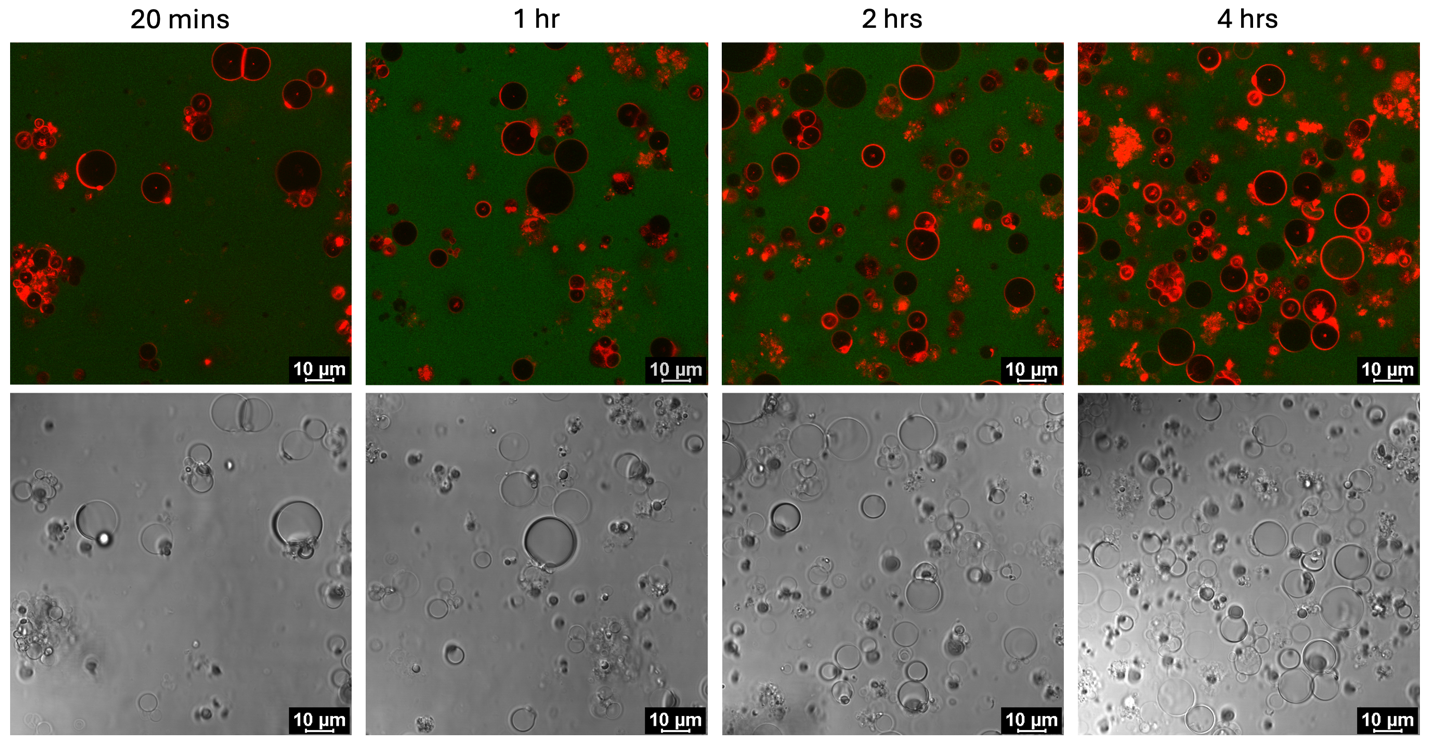

Figure S17**. Confocal microscopic images of fluorescein added to the POPC+OA-CSM over time.** We examined the fluorescein permeability over time (20 mins, and 1, 2, 4 hrs). We observed some green fluorescence intensity inside the membrane, which shows that fluorescein permeated into the CSMs, at longer times– particularly at 4 hours, where a few protocells had nearly the same fluorescence intensity inside as outside. However, none of the structures accumulated fluorescein in their interiors in the manner observed when the nonfluorescent substate, FDA was added (see Figure 6).

### References.
